# Natural naphthoquinones isolated from *Lithospermum erythrorhizon* suppress dextran sulfate sodium-induced murine experimental colitis

**DOI:** 10.1101/830224

**Authors:** Wenxue Sun, Hongwei Han, Zhaoyue Wang, Zhongling Wen, Minkai Yang, Yinsong Wang, Jiangyan Fu, Lu Feng, Xinhong Xu, Tongming Yin, Xiaoming Wang, Guihua Lu, Jinliang Qi, Hongyan Lin, Yonghua Yang

## Abstract

The purpose of this study was to explore the effects of natural shikonin and its derivatives on mice experimental colitis induced by dextran sulfate sodium, and to investigate the underlying mechanisms in *vivo*. Our results suggested that, intragastric administration of single compound like shikonin and its derivatives contributed to attenuating symptoms of malignant induced by DSS. Meanwhile, shikonin or its derivatives could also remarkably reduce the disease activity index and histopathological scores, suppress the levels of pro-inflammatory cytokines (including IL-6, IL-1β and TNF-α), while increase that of inflammatory cytokine IL-10 in serum. Additionally, both shikonin and alkanin were found to restrain the levels of COX-2, MPO and iNOS in serum and colonic tissues. Moreover, western blotting results demonstrated that shikonin and its derivatives could inhibit the activation of the NLRP3 inflammasome and the NF-κB signaling pathway, relieve the DSS-induced disruption of colonic epithelial tight junction (TJ) in colonic tissues. Further, docking simulation had been performed to prove that shikonin and its derivatives could bind to the active sites of NLRP3 inflammasome and the NF-κB to generate an effective inflammatory effect. Taken together, our experimental data can provide some evidence for the potential use of shikonin and its derivatives to treat the inflammatory bowel disease (IBD).

## Introduction

Inflammatory bowel disease (IBD), a kind of refractory and relapsing gastrointestinal inflammatory disease, has not only affected the living quality of patients, but increased the risk of colon cancer[1, 2]. Recently, an increasing amount of evidence indicates that the incidence of colorectal cancer (CRC) in IBD patients is generally much higher than that in general population[3, 4]. Currently, traditional treatment strategies for IBD can mainly be classified as three categories, namely, amino-salicylic acid agents, glucocorticoids and immunosuppressive agents[5]. Typically, 5-aminosalicylic acid remains the preferred drug to treat IBD, which can directly treat enteritis through inhibiting inflammatory reaction in the intestinal tract and exerting the antibacterial function. Nonetheless, it will be quickly absorbed by the small intestine if taken orally, leading to acute and chronic renal injury; besides, the drug can not reach the inflammatory site, which can hardly achieve the treatment goal[6]. In addition, glucocorticoid is likely to develop dependence after long-term application, and disease recurrence is common after drug withdrawal[7]. By contrast, immunosuppressant is suitable for patients with chronic enteritis, which can inhibit lymphocyte proliferation to achieve the therapeutic effect; nonetheless, it is associated with strong toxic side effects[8]. Fortunately, the emergence of anti-tumor necrosis factor (anti-TNF) therapy has made a major advance in the treatment of patients with IBD for the past two decades[9]. However, despite the therapy is useful in over half of IBD patients, a substantial proportion of patients are primary non-responders or lose response with long term use[10, 11]. What’s more, this therapy also exits some disturbing safety issues encompassing the increased risk of infections and malignant tumor in some groups[12, 13]. These factors pushed the need for finding some other safe, mild and durable effective drugs to cure IBD.

The disease characteristics of IBD and shortcomings of current treatments force us to turn to the traditional Chinese medicine (TCM). *Lithospermum erythrorhizon* is a commonly used TCM in the Chinese pharmacopoeia of the People’s Republic of China. Typically, it is abundant in shikonin derivatives, which have anti-tumor, anti-virus, and anti-inflammatory activities[14–18]. Numerous studies have indicated that shikonin and some of its derivatives are the main active ingredients in *Lithospermum erythrorhizon*[19–21]. There are dozens of species of shikonin in the Boraginaceae plants, among which, the representative ones are shikonin, *β, β*-dimethylacryl-shikonin, acetyl-shikonin, 5, 8-dihydroxy-1, 4-naphthoquinone, and alkanin (the enantiomeric of shikonin) with high contents[22, 23]. They belong to the naphthoquinones that are collectively known as shikonins. Typically, shikonin and its derivatives, which are isolated from the root of *Lithospermum erythrorhizon*, possess the advantages of bright color, long lasting effect, and non-toxic side effects. It has been reported that the crude extract of *Arnebia euchroma* is effective in rats with experimental colitis[24]. Whereas, the efficacy of each individual component of the crude extract has not been study detailly. In this work, we will further study and compare the effectiveness of some natural shikonins with high content in *Lithospermum erythrorhizon* for IBD treatment and to probe the potential mechanisms.

## Materials and Methods

### Chemicals, Regents and antibodies

Shikonin (SK, #B21682; CAS NO. 517-89-5; LOT: R10J8F39560), alkanin (AK, #B50783, CAS NO. 517-88-4; LOT: P05M7F14232), naphthoquinone (#AA18930, CAS NO. 475-38-7), acetyl-shikonin (acetyl-SK, #B21508, CAS NO. 24502-78-1; LOT: R21J7F16576) and *β*, *β*-dimethylacryl-shikonin (*β*, *β*-dimethylacryl-SK, #B24012, CAS NO. 24502-79-2; LOT: P12M7F11131) were ordered from Shanghai yuanye Bio-Technology Co., Ltd (Shanghai, China). All of them were characterized by ^13^C NMR, ^1^H NMR, melting properties and MS analysis, which were exactly accord with the structure described (***SI Appendix***). DSS (white powder; CAS NO. 9011-18-1; LOT: Q6182; M.W. = 36000 – 50000 Da) was purchased from MP Biomedicals (Shanghai, China). Mesalazine (#A600043-0050; CAS NO. 89-57-6; LOT: B326BA1390) was purchased from Sigma-Aldrich (Shanghai, China). Murine TNF-α (#ab208348), IL-1β (#ab197742), IL-6 (#ab100712), IL-10 (#ab108870) Elisa assay kits were purchased from Abcam (Cambridge, England). COX-2 (#SBJ-M0847), iNOS (#SBJ-M0041) and MPO (#SBJ-M0329) Elisa assay kits were ordered from Nanjing SenBeiJia Biological Technology Co., Ltd., (Nanjing, China). Primers were purchased from Genscript (Nanjing, China). All molecular docking results were in strict accordance with operation specification of Discovery Studio 3.5. The docking calculation of mesalazine, shikonin and its derivatives was showed in S1 Table.

PMSF (phenylmethanesulfonyl fluoride), RIPA lysis buffers (#P0013B) were purchased from Beyotime Institute of Biotechnology (Haimen, China). BCA protein assay kit (#23227) was purchased from Pierce (Rockford, IL, USA). anti-iNOS (#E1A8233A), anti-NF-κB p65 (#10745-1-AP) and anti-GAPDH (#10494-1-AP) were purchased from Enogene (Nanjing, China). Goat anti-mouse IgG (H+L), HRP conjugate (#SA00001-1) and Goat anti-rabbit IgG (H+L), HRP conjugate (#SA00001-2) were purchased from Proteintech (Wuhan, China). anti-NLRP3 (#AF2155), anti-STAT3 (#AF1492) were purchased from Beyotime Biotechnology (Nanjing, China). anti-ZO-1 (#WL03419), anti-Occludin (#WL01996), anti-Claudin-1 (#WL03073), anti-ASC (#WL02462), anti-IL-1β (#WL02257), anti-COX-2 (WL01750), anti-p-STAT3 (#WLP2412), anti-IKBα (#WL00148), anti-p-IKBα (#WL02495) and anti-p-NF-κB p65 (Ser536) (#WL02169) were purchased from Wanlei Biotech Co. Ltd (Shenyang, China), anti-VCAM-1 (#GB11336), anti-Caspase1 (#GB11383) and anti-β-actin (#GB11001) were purchased from Servicebio Technology Co. Ltd (Wuhan, China). anti-IKKα (#2682) and anti-IKKβ (#2678) were purchased from Cell Signaling Technology Inc. (Beverly, MA, USA). ECL Kit (#34077) was purchased from Thermo Scientific (USA). Haematoxylin-Eosin/HE Staining Kit (#20170804) was purchased from Solarbio Science & Technology Co. Ltd (Beijing, China).

### Animals

A total of 48 health male C57BL/6 mice weighted 18-20 g were selected for *in vivo* experiment. All of them were purchased from Model Animal Research Center of Nanjing University (Nanjing, China) and were randomly divided into eight groups for 1 week to let them adapt to their new environment, each consisting of six animals, including the model (the dextran sulfate sodium; 3.5% (w/v) DSS), SK (25 mg/kg), AK (25 mg/kg), naphthoquinone (25 mg/kg), acetyl-SK (25 mg/kg), *β*, *β*-dimethylacryl-SK (25 mg/kg), mesalazine (positive drug; 200 mg/kg) and control group. All mice housed under a 12 hours light-dark cycle at 22 ± 2°C and 60% ± 5% humidity. Animals were supplied with standard diet and sterile water ad libitum for 1 week, then all the groups mice except for control group were exposed to 3.5% DSS, which was dissolved in drinking water. The drugs were given orally once daily started at the same time as the DSS treatment and continued 1 week. All animal experiments and welfare were treated in strict accordance with the relevant Guidelines for Care and Use of Laboratory Animals of Nanjing University and approved by the Animal Ethics Committee of the Ministry of Science and Technology of China (2006). All efforts were aimed at reducing the number and suffering of experimental animals while meeting the needs of the experiment.

### Establishment of acute enteritis model and treatment with SK and its derivatives

Acute colitis was induced by free intake of sterile water containing 3.5% DSS for one week. Another six groups (n=6 mice per group) of mice which also fed orally with 3.5% DSS in sterile water accompanied with the usage of SK, AK, naphthoquinone, acetyl-SK, *β*, *β*-dimethylacryl-SK and mesalazine (the positive control group) every day. All drugs were dissolved in olive oil and were perfused 200 μL per mice at a predetermined dose by gavage. In the DSS group, the mice fed orally with the same amount of olive oil. In the blank control group, the mice received only sterile water and fed orally with the same amount of olive oil. There was no significant difference in water intake among all groups. Body weight and the disease activity index (DAI) were measured and recorded everyday[25]. The DAI was calculated according to the standard listed in S2 Table. After one week of treatment, all the mice monocular eyeball was removed for blood collection under lightly anesthetized with ether. Then, the mice were euthanized by cervical dislocation. The colonic tissues were removed for pathological analysis. The same protocol was carried out at least three times in independent experiments.

### Measurement of Cytokines, COX-2, MPO and iNOS in serum

The expressions of various cytokines (IL-6, IL-1β, TNF-α and IL-10), COX-2, MPO and iNOS in the serum of mice were detected by an enzyme-linked immunosorbent assay kit according to the product’s guidelines.

### Haematoxylin & Eosin (HE) staining

Small sections of colonic colonic tissue were fixed in 10% buffered formalin and embedded in paraffin. Then, sections were stained with haematoxylin and eosin.

Histologic evolution of colonic mucosa was carried out in a blinded fashion by a pathologist which has been widely used as evaluation criterion[24].

### Immunohistochemical analysis

Immunohistochemistry was performed for Cyclooxygenase-2 (COX-2), NF-κB, NLRP3. The tissues were incubated with primary antibody at 4 °C for at least 12 h after deparaffinized and rehydrated. The Alexa Fluor 488 labeled anti-mouse secondary antibody (Invitrogen, USA) were treated 60 min at 25 °C. Signals were developed with Hematoxylin and DAB (Dako, Agilent Technologies, USA).

### Western bolt analysis

Colonic samples were lysed with RIPA lysis buffer which containing 1% PMSF. The harvested protein concentrations were assessed by using the BCA Protein Assay kit. 60-80 µg of total protein per sample were loaded onto 10 % SDS PAGE and transferred onto PVDF membranes for immunoblot analysis[25]. The blots were detected by super ECL detection reagent (Yeasen Biotech Co., Ltd) according to the schemes of the manufacturer and then exposed to film by digital camera (Tanon 5200, Tanon, China). Finally, the immunoblots were quantified by densitometry with Image J Software (National Institutes of Health, BetheSEMa, Maryland, USA), and β-actin or GAPDH was used as loading control.

### RNA isolation and quantitative real-time PCR analysis

The extraction of colonic tissues total RNA was according to the manufacture instructions by using RNAprep pure Tissue Kit (TIANGEN BIOTECH, BeiJing, China). After that, the first step product was converted to cDNA using FastKing RT Kit (With gDNase) (TIANGEN BIOTECH, BeiJing, China). Then, the collected cDNA solution was used for qRT-PCR using Power SYBR® Green PCR Master Mix (Roche, Indianapolis, USA) according to the schemes of the manufacturer, which was carried out by the BioRad CFX96 TouchTM RealTime PCR Detection System (BioRad, CA). The sequences of each primer are listed in S3 Table.

### Docking simulation

Molecular docking of the compounds binding to the three-dimensional X-ray structure of MPO (PDB code: 4C1M), NF-κB (PDB code: 4DN5), NLRP3 (PDB code: 6NPY), TNF-α (PDB code: 2AZ5), IL-1β (PDB code: 3O4O), IL-6 (PDB code: 5U98), IFNγ (PDB code: 3OQ3) and IL-10 (PDB code: 5T5W) were performed using Discovery Studio (version 3.5). SK, AK, naphthoquinone, acetyl-SK, *β*, *β*-dimethylacryl-SK and mesalazine were constructed, minimized and prepared. The RCSB Protein Data Bank (http://www.rcsb.org/pdb/home/home.do) provides the crystal structures of the protein complex download. All bound waters and ligands were removed from the protein. After the molecular docking, Discovery Studio (version 4.5) were used for analysis of types of interactions between the ligands and docked proteins.

### Data and Statistical analysis

Results were presented as mean ± S.E.M. of three independent experiments. Statistical comparisons between the treated and untreated groups were performed using one-way ANOVA using GraphPad PRISM5 (Graphpad Inc., La Jolla, USA). *P* < 0.05 was considered to be significant; *P* < 0.01 were considered to be very significant.

## Results

### SK and its derivatives protected the colon from DSS-induced damage in an acute ulcerative colitics mouse model

To investigate the effects of SK and its derivatives on colitis, we established an acute DSS-induced ulcerative colitis mouse model. In the model, mice were given either sterile water or 3.5% DSS dissolved in sterile water. Six other groups received 3.5% DSS and were administered SK, naphthoquinone, *β*, *β*-dimethylacryl-SK, acetyl-SK, AK, mesalazine orally both at the beginning of the experiment (n=6 mice per group). At the end of this experiments, SK and its derivatives could attenuate the severity level of inflammatory significantly. In the case of the colorectum length, DSS group mice suffered a shortening of about 30% when compared to normal mice. While SK and its derivatives significantly prevented this shortening (Fig 1A and 1B). Moreover, SK and its derivatives treated mice showed improvements in body weight, disease activity index (DAI) and alleviation of splenomegaly (Fig 1C-1F). SK and its derivatives thus ameliorated the colon syndromes of DSS-induced damage in mice.

**Fig 1.**
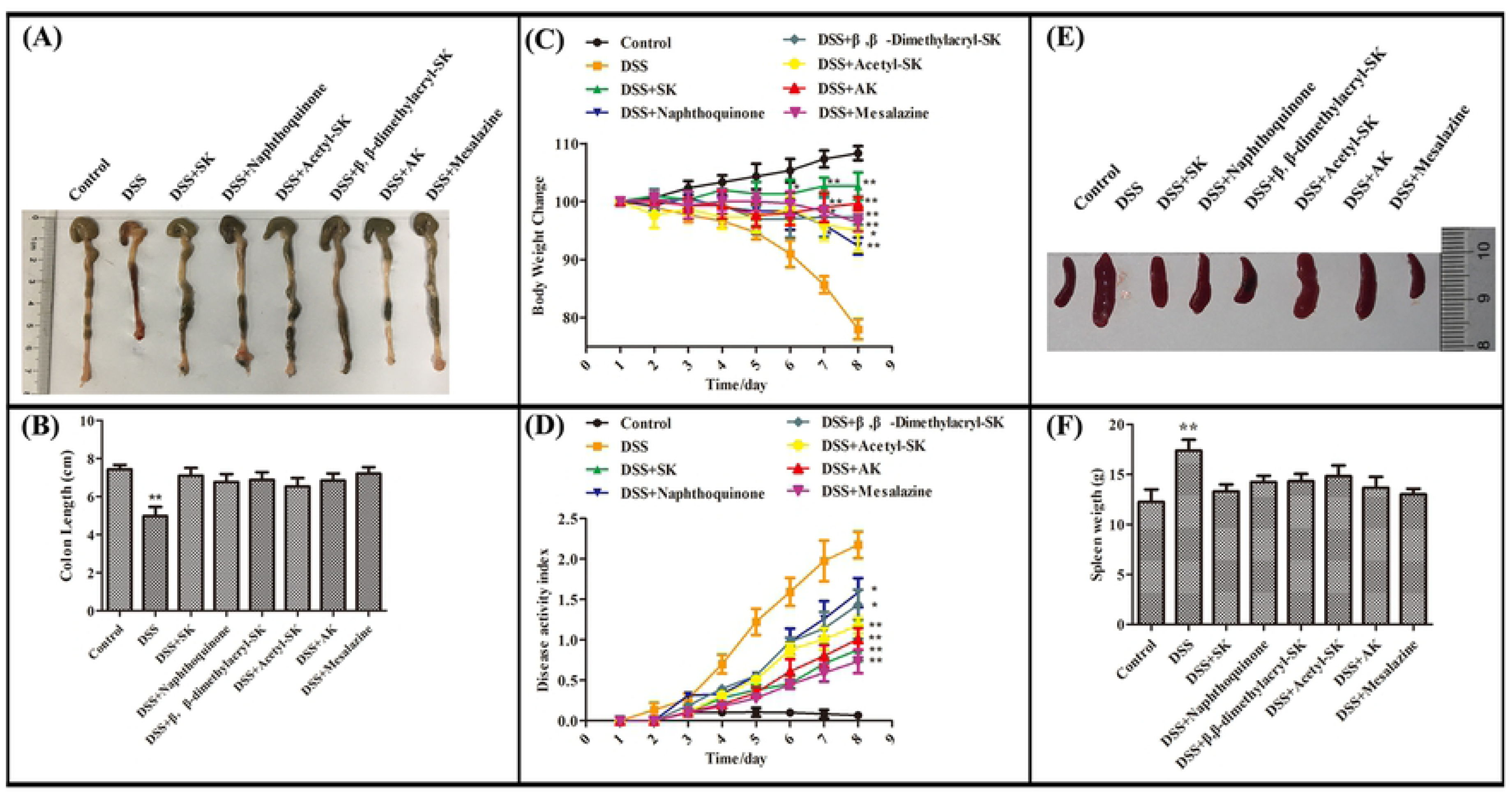
SK and its derivatives protect the colon from DSS-induced damage. **(A)** Macroscopic observation of colon length. **(B)** Bar graph showing colon length. \*\**p*<0.01 versus control group. **(C)** Pattern of daily weight changes. \**p*<0.05, \*\**p*<0.01 versus DSS-treated group. **(D)** Disease activity index of mice in each group. \**p*<0.05, \*\**p*<0.01 versus DSS-treated group. **(E)** Macroscopic observation of spleen size. **(F)** Bar graph showing spleen weight. Data are shown as mean ± S.E.M, \**p*<0.05 versus control group (n=6 per group).

### SK and its derivatives limited the expression of pro-inflammatory cytokines

Once the circulating leukocytes were recruits into the colon, it will lead to the release of pro-inflammatory mediators, including pro-inflammatory cytokines such as IL-6, IL-1β, TNF-α and pro-inflammatory synthases such as inducible nitric oxide synthase (iNOS) and cyclooxygenase-2 (COX-2). To determine whether SK and its derivatives attenuates the release of pro-inflammatory cytokines, therefore protecting the intestinal mucosa, we detected the levels of IL-6, IL-1β, TNF-α and IL-10 (anti-inflammatory cytokine) in the supernatant of cultured colons of 3.5% DSS-exposed mice given control diet or SK and its derivatives. As shown in Fig 2A-2D, treatment with SK and its derivatives at a dosage of 25 mg/kg, significantly reduced the levels of pro-inflammatory cytokines (IL-6, IL-1β and TNF-α) and added the level of anti-inflammatory cytokine (IL-10) in the serum. Among them, SK showed the best anti-inflammation effect that was similar to mesalazine group at a dosage of 200 mg/kg. As shown in Fig 2E-2H, SK and mesalazine (the positive control) were found to reduce the levels of pro-inflammatory cytokines (IL-6, IL-1β and TNF-α) and increased the level of anti-inflammatory cytokine (IL-10) in a dose-dependent manner and SK treatment resulted in better anti-inflammatory effect at a dose of 25 mg/kg. The above results suggest that SK and its derivatives may improve the severity of colon damage induced by DSS through inhibiting the expression of pro-inflammatory cytokines.

**Fig 2.**
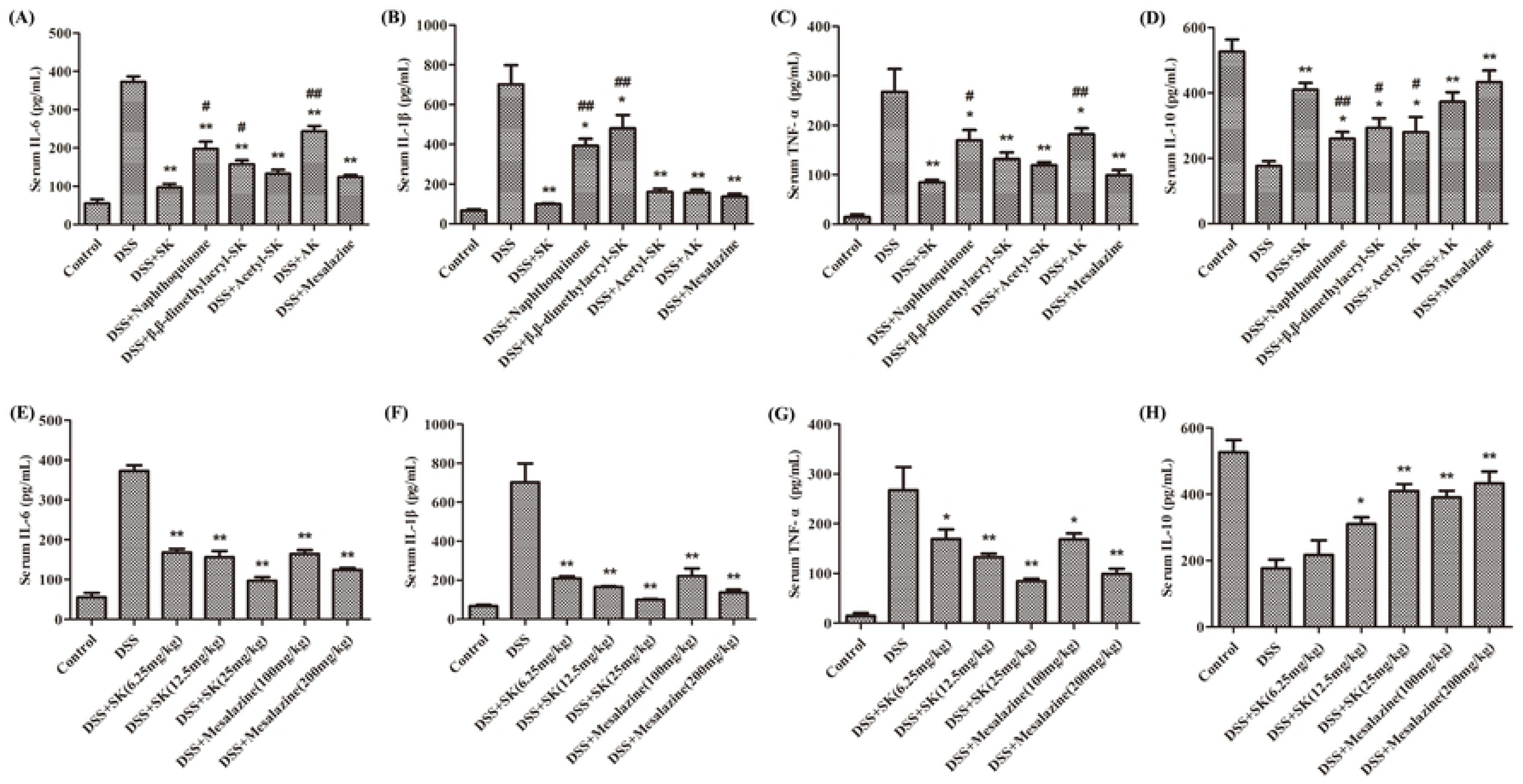
SK and its derivatives exhibited a stronger ability in decreasing the amount of pro-inflammatory cytokines and increasing the level of anti-inflammatory cytokine in serum. **(A-D)** The protein level of cytokines IL-6, IL-1β, TNF-α and IL-10 in serum were determined by ELISA (DSS: 3.5%; SK, Naphthoquinone, β, β-dimethylacryl-SK, Acetyl-SK, AK: 25 mg/kg; Mesalazine: 200 mg/kg). **(E-H)** Effects of different concentrations of SK on the levels of cytokines IL-6, IL-1β, TNF-α and IL-10. Data are shown as mean ± S.E.M, \**p*<0.05, \*\**p*<0.01 versus control group; #*p*<0.05, ##*p*<0.01 versus DSS-treated group (n=6 per group).

### SK and its derivatives strengthened the prevention of colon from DSS-induced the histological damage

The severity level of colon tissues was assessed by distortion of crypts, loss of goblet cells, infiltration of inflammatory cells, and severe mucosal damage which were observed by HE staining. The results shown in Fig 3A and 3B indicated that the serious deterioriation of the colons were markedly protected by SK and its derivatives. In contrast, we found SK and AK showed better effects than other derivatives and the protection of them are similar to that of mesalazine.

**Fig 3.**
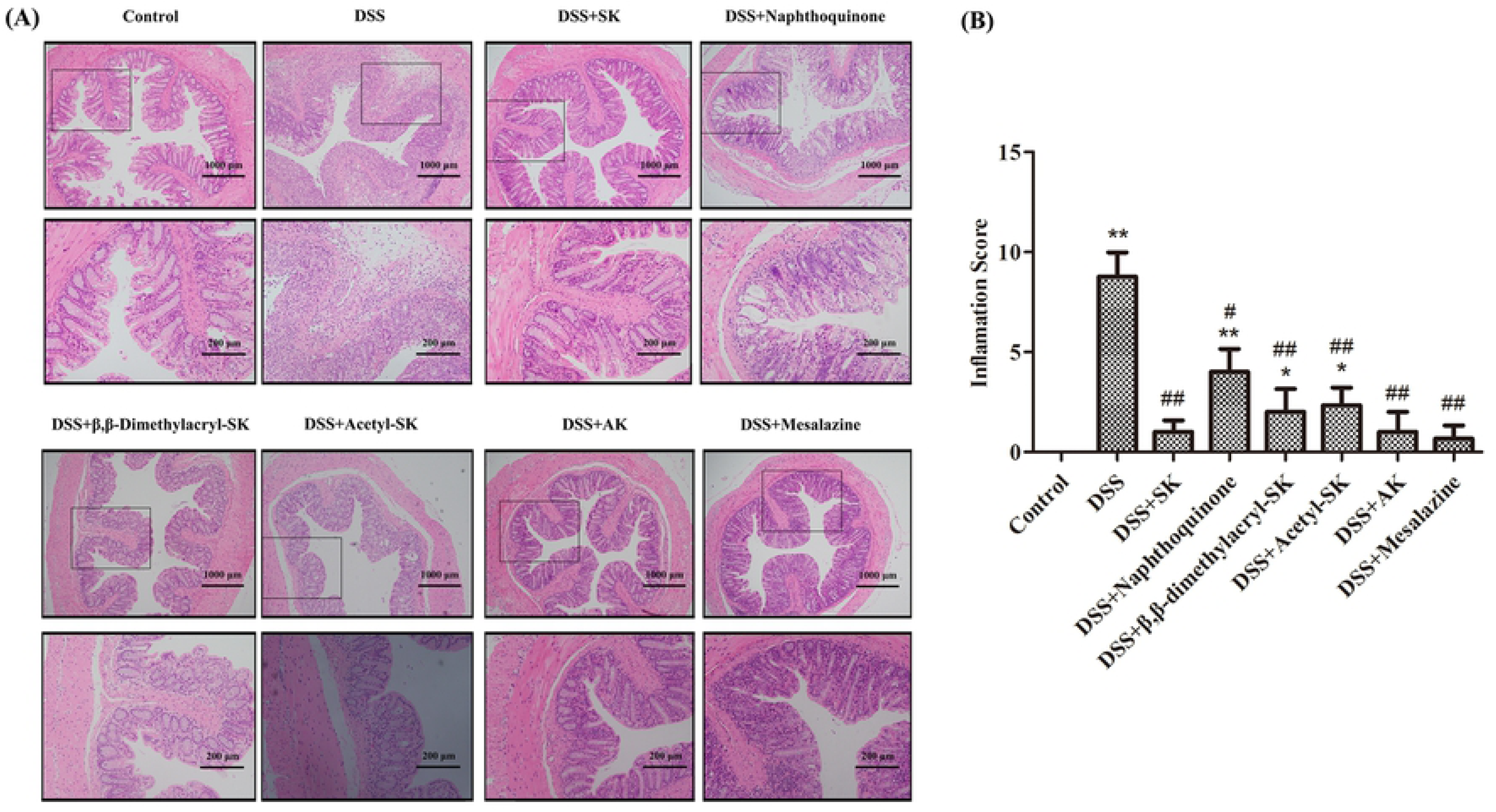
Colon histology and inflammation score of mice in each group (H&E staining 200× and 400×). **(A)** The colon histology sections were stained with hematoxylin and eosin. The colon histology sections of DSS-treated group mice revealed sever pathology with observation of submucosal edema with extensive inflammatory cell infiltration. The frame indicates the region magnified in the bottom panel. **(B)** Inflammation scores of each group. Data are shown as mean ± S.E.M, \**p*<0.05, \*\**p*<0.01 versus control group; #*p*<0.05, ##*p*<0.01 versus DSS-treated group (n=6 per group).

### SK and its derivatives reduced the activities of MPO, COX-2 and iNOS in serum

MPO is an enzyme which is present largely in neutrophils, as well as in monocytes and macrophages in small concentrations. The concentrations of MPO, COX-2 and iNOS in all the groups of mice are shown in Fig 4. The MPO activity is an indicator of neutrophil infiltration in inflamed colon tissues[26]. Administration of DSS increased the activity of MPO as compared to the normal mice. SK and AK administration markedly reduced the concentrations of MPO, COX-2 and iNOS as compared to DSS-induced mice.

**Fig 4.**
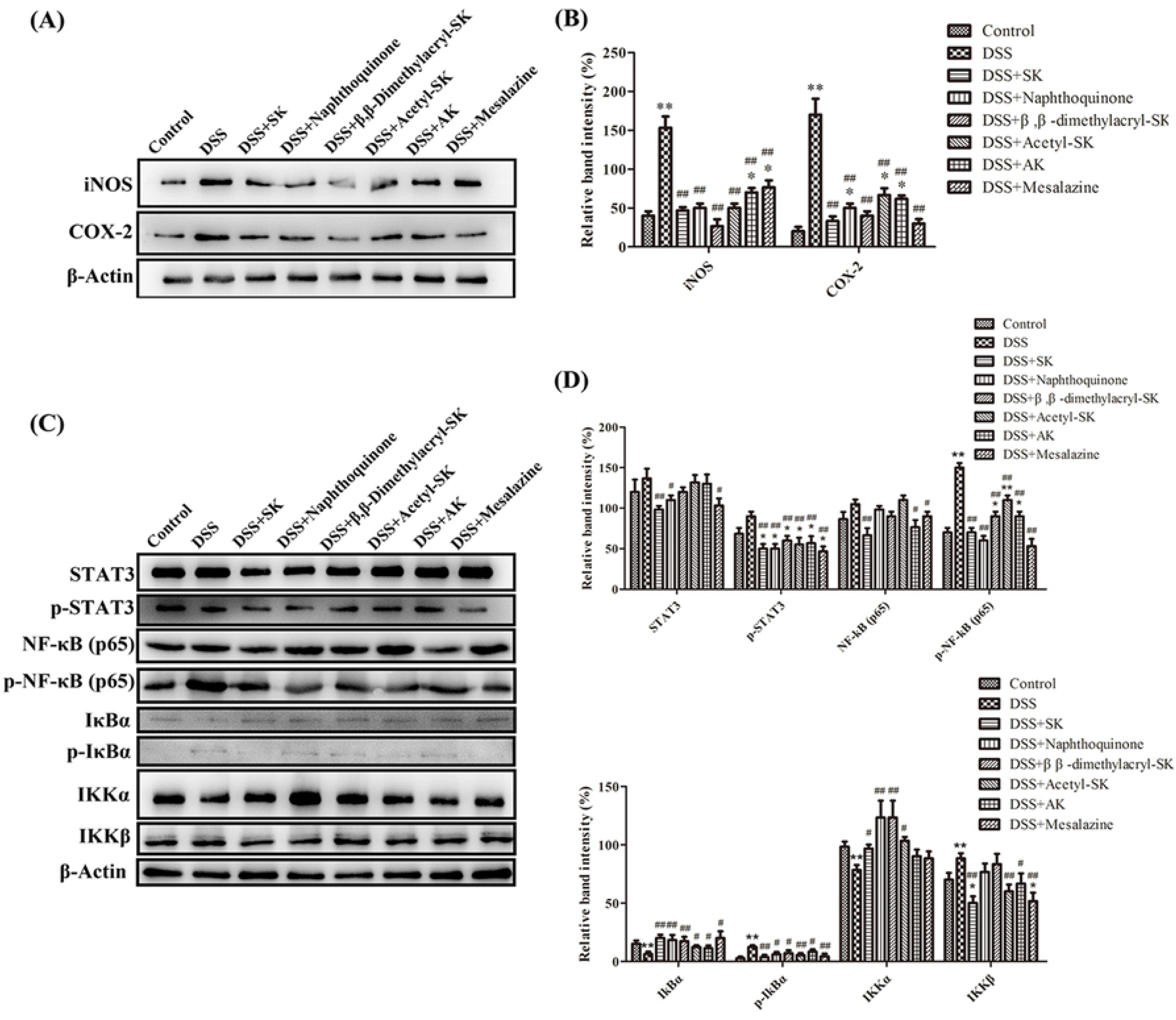
SK and its derivatives reduced the level of COX-2, MPO and iNOS in the serum of mice. The concentrations of COX-2 **(A)**, MPO **(B)** and iNOS **(C)** were measured using an enzyme-linked immunosorbent assay kit. Each point represents the mean ± S.E.M. from three replicates (*, *P* < 0.05; **, *P* <0.01 versus control; #, *P* < 0.05; ##, *P* <0.01 versus DSS group).

### SK and its derivatives regulate the expression of COX-2, iNOS and the activation of NF-κB in colonic tissues

Transcription factors of the NF-κB family play a vital role in the regulation of genes, which can influence the immune and inflammatory response[27]. Many evidences have verified that NF-κB plays an important role in the development of inflammatory bowel disease. In his disease, NF-κB activation is observed in inflamed colonic mucosa, in which it leads the production of COX-2 and iNOS. To evaluate the mechanism of SK, we detected the phosphorylation level of STAT-3, NF-κB p65 and IκB. The outcome shown in Fig 5 suggested that the phosphorylation levels of STAT-3, NF-κB p65 and IκB were markedly up-regulated in DSS-treated group compared with the control group. While SK and its derivatives can effectively inhibit the activation of NF-κB and STAT-3, thus decreasing the generation of COX-2 and iNOS in the colon homogenates.

**Fig 5.**
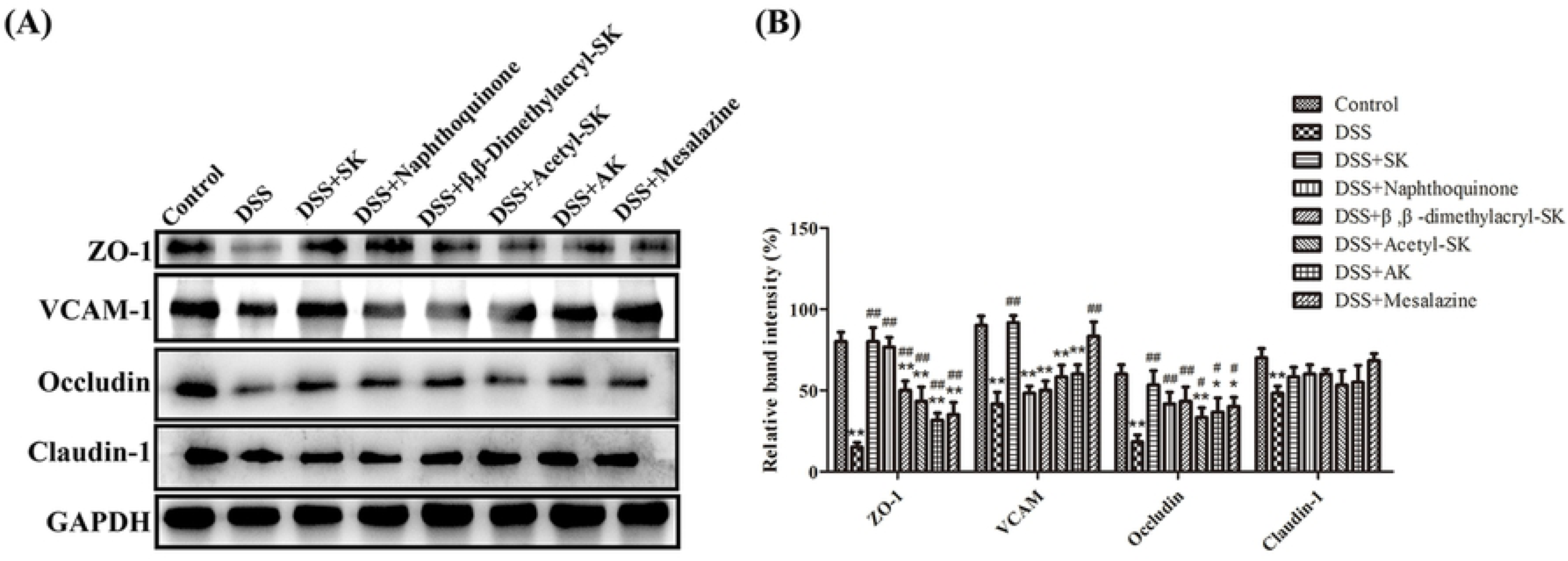
Effect of SK and its derivatives on COX-2, iNOS expression, and NF-κB signal pathway in DSS-induced acute colitis. **(A)** Western blot analysis of COX-2 and iNOS levels in colonic tissues in each group. Data shown are representative of three independent experiments; **(B)** Relative protein expression ratios of COX-2 and iNOS were determined by densitometry; **(C)** Western blot analysis of proteins expression level of NF-κB signal pathway; **(D)** Relative protein expression ratios of were determined by densitometry. Each point represents the mean ± S.E.M. from three replicates (*, *P* < 0.05; **, *P* <0.01 versus control; #, *P* < 0.05; ##, *P* <0.01 versus DSS group).

### SK and its derivatives relieved DSS-induced colonic epithelial tight junction disruption in mice

ZO-1, VCAM-1, Occludin and Claudin-1 proteins play an important role in epithelial tight junction, which can protect mucosa epithelial cells from breaching by the harmful substances, maintaining cellular permeability and integrity, and ensuring homeostasis of internal environment[28]. Therefore, we further investigated the effect of SK and its derivatives on epithelial tight junction proteins. The results suggested that the expression of ZO-1, VCAM-1, Occludin and Claudin-1 were remarkably reduced in DSS group compared to the control group. However, SK and its derivatives can restore the expression of these epithelial tight junction proteins (Fig 6).

**Fig 6.**
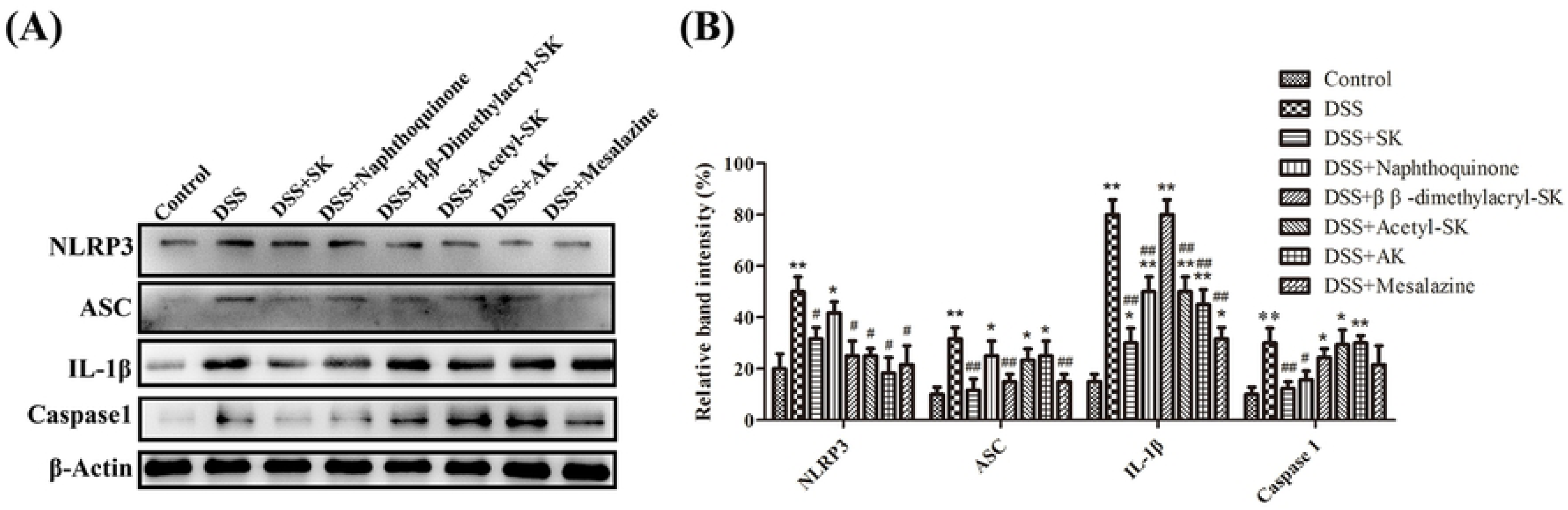
SK and its derivatives relieved DSS-induced colonic epithelial tight junction disruption in mice. **(A)** Protein expression of ZO-1, VCAM-1, Occludin and Claudin-1 in the colon tissues were analyzed by Western blot. **(B)** Relative protein expression ratios of were determined by densitometry and normalized to GAPDH. Each point represents the mean ± S.E.M. from three replicates (*, *P* < 0.05; **, *P* <0.01 versus control; #, *P* < 0.05; ##, *P* <0.01 versus DSS group).

### SK and its derivatives inhibit the activation of NLRP3 inflammasome in colonic tissues

Inflammasomes, a multiprotein oligomer, have been widely reported that it can promote the activation of CASP1/caspase 1 and maturation of proinflammatory cytokine IL1B (interleukin 1, β)[29]. Many researches have revealed that NLRP3 affects several inflammatory disorders to a large extent including colitis. As shown in Fig 7, NLRP3, ASC, caspase-1 and IL-1β expressions in DSS group were significantly increased when compared to the control group. However, these changes were receded by SK and its derivatives with different degrees.

**Fig 7.**
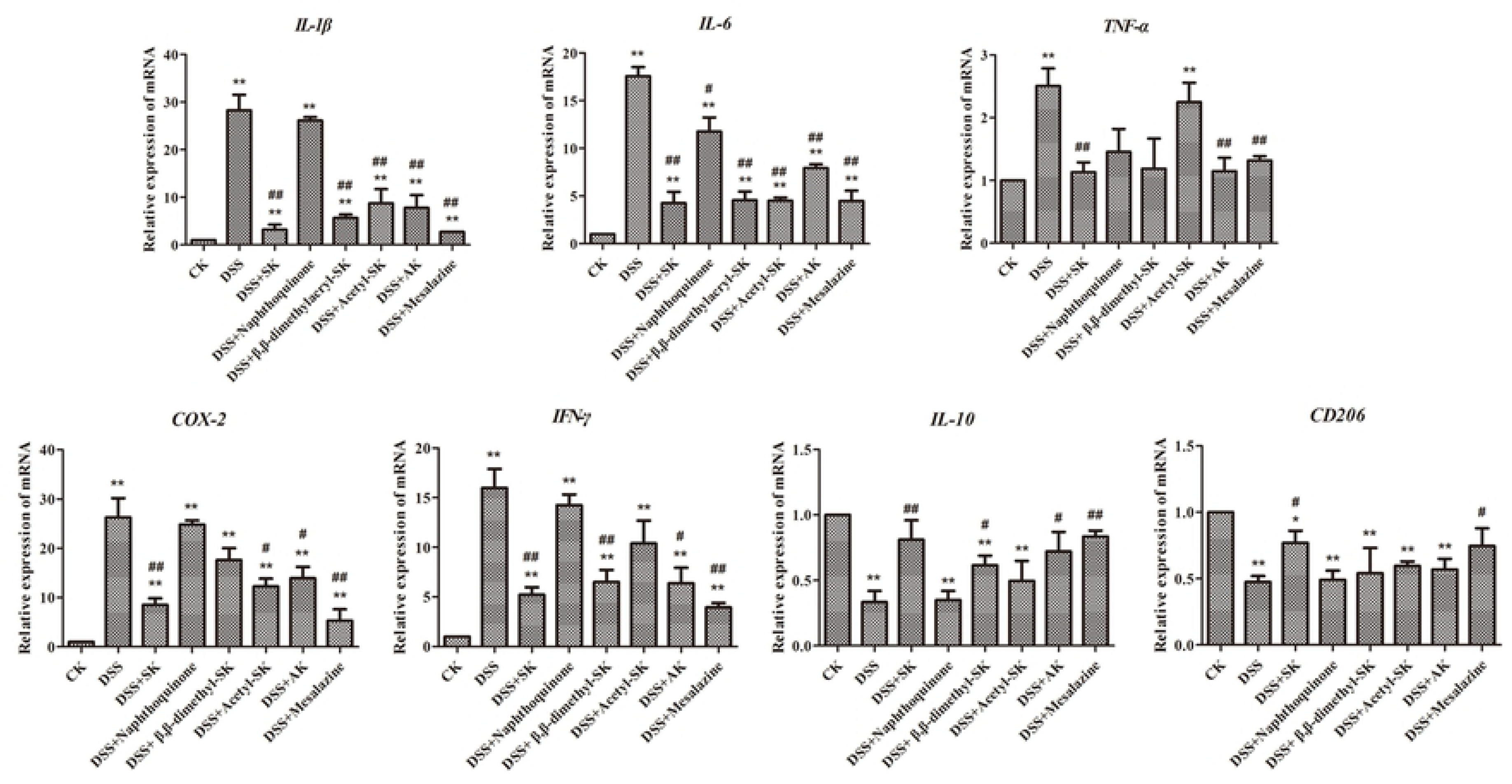
SK and its derivatives reduced the level of NLRP3 inflammasome activation in colonic tissues. **(A)** Protein levels of NLRP3, ASC, caspase-1, and IL-1*β* were determined by Western blotting. *β*-Actin was used as a control. **(B)** Relative protein expression ratios of were determined by densitometry and normalized to *β*-Actin. Each point represents the mean ± S.E.M. from three replicates (*, *P* < 0.05; **, *P* <0.01 versus control; #, *P* < 0.05; ##, *P* <0.01 versus DSS group).

### SK and its derivatives reduced the mRNA level of cytokines and protease

Real time-PCR analysis and mRNA expressions of IL-1β, IL-6, TNF-α, IFNγ, COX-2, IL-10 and CD206 were also detected and the results were shown in Fig 8. The DSS treatment apparently enhanced mRNA levels of IL-1β, IL-6, TNF-α, IFNγ and COX-2 as compared to the control group, while SK and its derivatives can reduced the mRNA levels of these pro-inflammatory cytokines. On the contrary, the mRNA levels of anti-inflammatory cytokines (IL-10 and CD206) in the DSS treatment mice were lower than the control group. The subsequent SK administration increased IL-10 and CD206 mRNA effectively.

**Fig 8.**
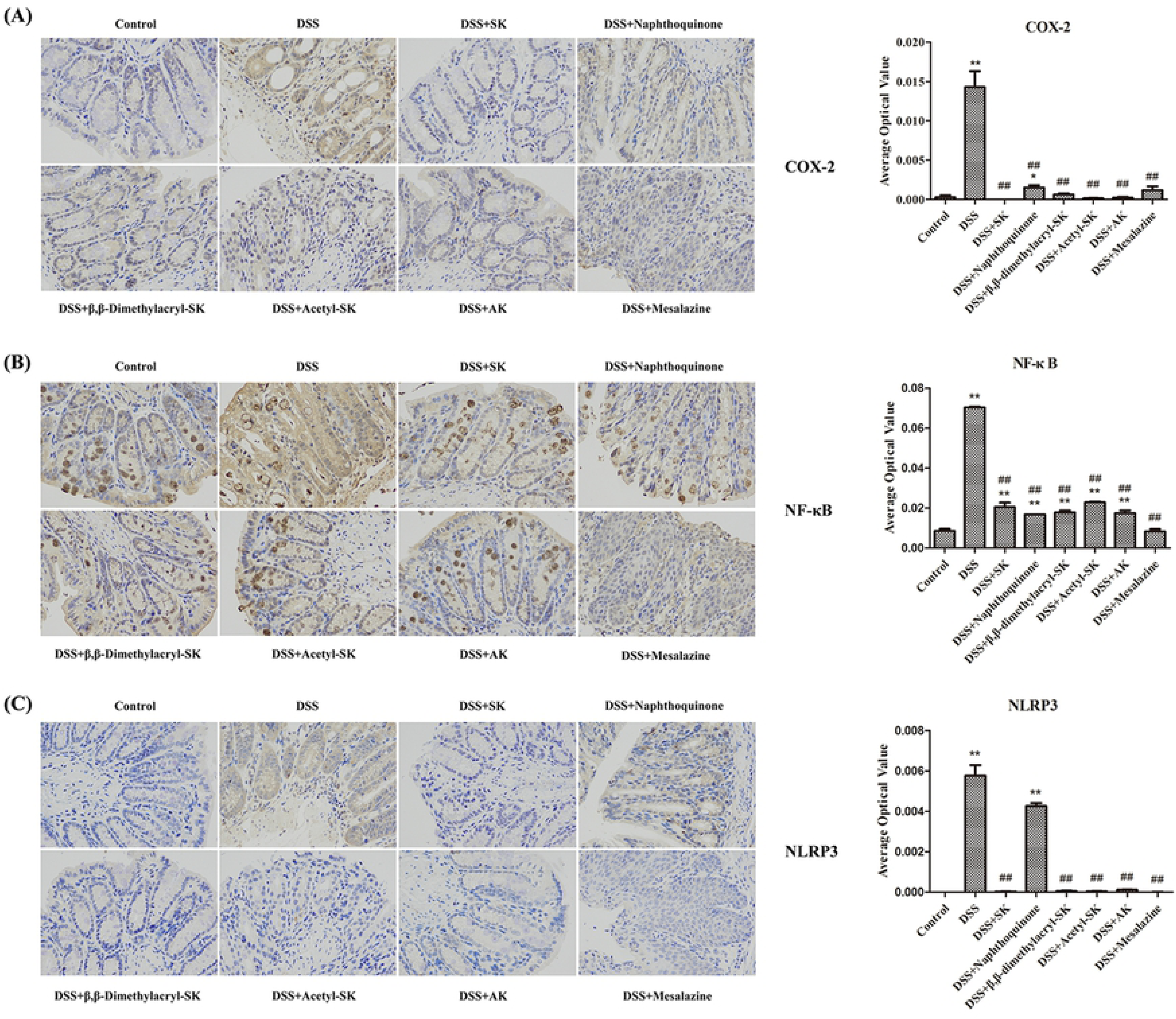
Real-time PCR quantification of signature genes in inflammation pathways in colon tissues of all groups. Down-regulation (%) was compared with vector control. Images are representative of three independent experiments. Data are mean ± S.E.M. of three independent experiments (*, *P* < 0.05; **, *P* <0.01 versus control; #, *P* < 0.05; ##, *P* <0.01 versus DSS group).

### SK and its derivatives attenuate the production of COX-2, NF-κB and NLRP3 in colonic tissues

Immuno-histochemical expressions of COX-2, NF-κB and NLRP3 in all the groups of mice are shown in Fig 9. The results demonstrated that DSS administration resulted in increased production of COX-2, NF-κB and NLRP3 in colonic tissues compared to the control group. Usage of SK and its derivatives were able to reduce the expression of COX-2, NF-κB and NLRP3 significantly (p < 0.05) as compared to mice induced with DSS.

**Fig 9.** SK and its derivatives attenuate the expression of COX-2, NF-κB and NLRP3. **(A)** Immuno-histochemical analysis of COX-2 in each group. **(B)** Immuno-histochemical analysis of NF-κB in each group. **(C)** Immuno-histochemical analysis of NLRP3 in each group. The slides were incubated with primary antibody. After the secondary antibody incubation, the slides were developed with DAB and counter-stained with hematoxylin. The images were taken at 400×magnification. Data are presented as means ± S.E.M. from three replicates (*, *P* < 0.05; **, *P* <0.01 versus control; #, P < 0.05; ##, P <0.01 versus DSS group).

### Molecular docking

Docking study has confirmed that SK and its derivatives anchored in the binding pocket of the protein perfectly through a series of binding mode, which may play a vital role in the conformation with them. As shown in. S1-S4 Figs, the binding mode of the compounds with proteins were stabilized mainly through hydrogen bond that are shown in the 2D and 3D diagrams. The binding energy of the compounds with proteins implies their anti-inflammatory activity. The whole results showed that the docking model had certain credibility and demonstrated that shikonin and its derivatives were potential anti-inflammatory agents compared to the positive drug mesalazine.

## Discussion

*Lithospermum erythrorhizon* is a kind of TCM, which has long been used for wound healing in ancient times. Typically, its roots are rich in a variety of bioactive substances, including natural naphthoquinones, which represents a continuing source to mine the therapeutic agents[30–32]. Among the naphthoquinones isolated from *Lithospermum erythrorhizon*, shikonin has attracted extensive interest due to its multiple biological activities, such as anti-inflammatory[18, 33], wound healing[34], anti-microbial[35], anti-ulcer[36], anti-thrombus[37] and anti-cancer activities[38, 39]. Hitherto, nearly 20 kinds of natural shikonin derivatives have been identified.

The anti-inflammatory effects of natural naphthoquinones have been extensively investigated. For instance, Fan *et al*. had investigated the effectiveness of hydroxynaphthoquinone mixture (HM) isolated from *Arnebia euchroma* on rats with experimental colitis induced by 2, 4, 6-tri-nitrobenzene sulfonic acid (TNBS) in 2013.[24] Their results suggested that HM could remarkably attenuate the clinical and histopathological severity of the TNBS-induced colitis in a dose-dependent manner. In addition, it could also alleviate loss of body weight, hematochezia and inflammation, reduce the macroscopic damage score, and improve the histological signs; moreover, it can evidently reduce the inflammatory infiltration, ulcer area and the severity of goblet cell depletion. Notably, HM, which is isolated from *Arnebia euchroma*, contains seven alkannin derivatives, including alkannin, deoxyalkannin, acetylalkannin, *β*, *β*-dimethylacrylalkannin, *α*-methylalkannin, isovalerylalkannin and *β*-acetoxyisovalerylalkannin. In 2012, Isabel *et al*. had reported that the oral administration of shikonin could attenuate the DSS-induced acute ulcerative colitis by blocking the activation of two major targets NF-κB and STAT-3[27]. However, the effectiveness of the commonly used shikonin and its natural derivatives on acute colitis has not been systematically investigated yet.

In this study, five representative natural naphthoquinones, namely, 5, 8-dihydroxy-1, 4-naphthoquinone, shikonin, acetyl-shikonin, *β*, *β*-dimethylacryl-shikonin and alkanin, had been selected to treat DSS-induced colitis in mice. The above results suggested that, the administration of these five natural naphthoquinones could ameliorate the clinical severity of the wasting disease, prevent the shortening the length of colorectum, and decrease the loss of body weight, improve the appearance of feces, and prevent bloody excrements. Such beneficial effects were further proved by histological evaluation, as evidenced by the markedly reduced severity and extent of inflamed tissue damage and infiltration of inflammatory cells. Typically, the relative expression of pro-inflammatory cytokines, including IL-6, IL-1*β* and TNF-α in serum, as well as their mRNA levels in colon tissues, suggested that these five natural naphthoquinones had beneficial effects. As mentioned earlier, MPO is a marker of neutrophil infiltration[26], which has been observed to be activated in several experimental colitis models, including the DSS-induced colitis. Therefore, MPO could be recognized as an indicator to quantify intestinal inflammation and evaluate the severity of inflammation[26]. In addition, the pro-inflammatory synthases, including iNOS and COX-2, are the vital inflammatory mediators with vital functions during the pathogenesis of ulcerative colitis[40]. The five natural naphthoquinones could suppress the MPO, iNOS and COX-2 activities, and such results were consistent with those found in histological examination, in which the inflammatory cell infiltration extents in colonic tissues were less than that in shikonin- and alkannin-treated animals than in 5, 8-dihydroxy-1, 4-naphthoquinone-, acetyl-shikonin- and *β*, *β*-dimethylacryl-shikonin-treated mice. Besides, results of Western boltting had further verified that shikonin could relieve the colitis in mice through inhibiting the activation of NF-κB and NLRP3 inflammasome, as well as the disruption of epithelial tight junction proteins (ZO-1, VCAM-1, Occludin and Claudin-1). Collectively, our work has provided supporting evidence for the clinical application of natural naphthoquinones, especially for shikonin.

## Conclusions

To sum up, through in *vitro* and in *vivo* experiments, we found that naphthoquinone compounds shikonin and some of its derivatives which were isolated from the root of the Chinese herbal medicine *Lithospermum erythrorhizon* could effectively attenuate the severity of DSS-induced colitis. This effect may due to the reduction of pro-inflammatory cytokines release, including IL-6, IL-1*β* and TNF-α in serum, as well as their mRNA levels in colon tissues, while the expression of anti-inflammatory cytokines (IL-10 and CD206) were increased. Moreover, they could suppress the MPO, iNOS and COX-2 activities, and such results were consistent with those observed in histological examination. Furthermore, changes in the expression of inflammatory regulatory proteins have been detected. These aforementioned results offer a kind of perspective for the potential use of shikonin and its derivatives to treat the IBD.

## Abbreviations

DSS: dextran sulfate sodium
IL-1β: interleukin1β
IL-6: interleukin6
TNF-α: tumor necrosis factor-α
IL-10: interleukin10
COX-2: cyclooxygenase-2
MPO: myeloperoxidase
iNOS: inducible nitric oxide synthase
ASC: apoptosis-associated speck-like protein containing CARD
NLRP3: nod-like receptor protein3
NF-κB: nuclear factor-kappa B
TJ: tight junction
IBD: inflammatory bowel disease
CRC: colorectal cancer
TCM: traditional Chinese medicine
SK: shikonin
AK: alkanin
acetyl-SK: acetyl-shikonin
*β*: *β*-dimethylacryl-SK, *β*-dimethylacryl-shikonin
DAI: disease activity index
HE: haematoxylin & Eosin
RIPA: radio immunoprecipitation assay
PVDF: polyvinylidene fluoride
HRP: horseradish peroxidase
UC: ulcerative colitis
ZO-1: zonula occludens
VCAM-1: vascular cell adhesion protein 1
HM: hydroxynaphthoquinone mixture
TNBS: 2, 4, 6-tri-nitrobenzene sulfonic acid

## Acknowledgements

This research was supported by the National Natural Science Foundation of China (NSFC) (21702100, 31771413, 31670298, 21907051), the Program for Changjiang Scholars and Innovative Research Team in University (IRT_14R27), and the Fundamental Research Funds for the Central Universities (020814380002, 020814380057).

## Author contributions

**Conceptualization:** Yonghua Yang, Hongyan Lin, Guihua Lu, Jinliang Qi.

**Data curation:** Zhongling Wen, Yinsong Wang, Jiangyan Fu, Lu Feng, Xinhong Xu, Tongming Yin.

**Formal analysis:** Zhongling Wen, Yinsong Wang, Jiangyan Fu. **Investigation:** Yonghua Yang, Hongyan Lin, Hongwei Han, Wenxue Sun. **Methodology:** Hongyan Lin, Xiaoming Wang, Guihua Lu, Jinliang Qi.

**Project administration:** Wenxue Sun, Hongwei Han, Zhaoyue Wang, Minkai Yang.

**Resources:** Xiaoming Wang, Guihua Lu, Jinliang Qi.

**Supervision:** Yonghua Yang, Hongyan Lin, Guihua Lu, Jinliang Qi.

**Validation:** Yinsong Wang, Jiangyan Fu, Yonghua Yang, Hongyan Lin, Guihua Lu, Jinliang Qi.

**Visualization:** Hongwei Han, Hongyan Lin.

**Writing – original draft:** Wenxue Sun, Hongyan Lin.

**Writing – review & editing:** Yonghua Yang, Hongyan Lin, Guihua Lu, Jinliang Qi.

## Supplementary Contents

**Sl Table.**
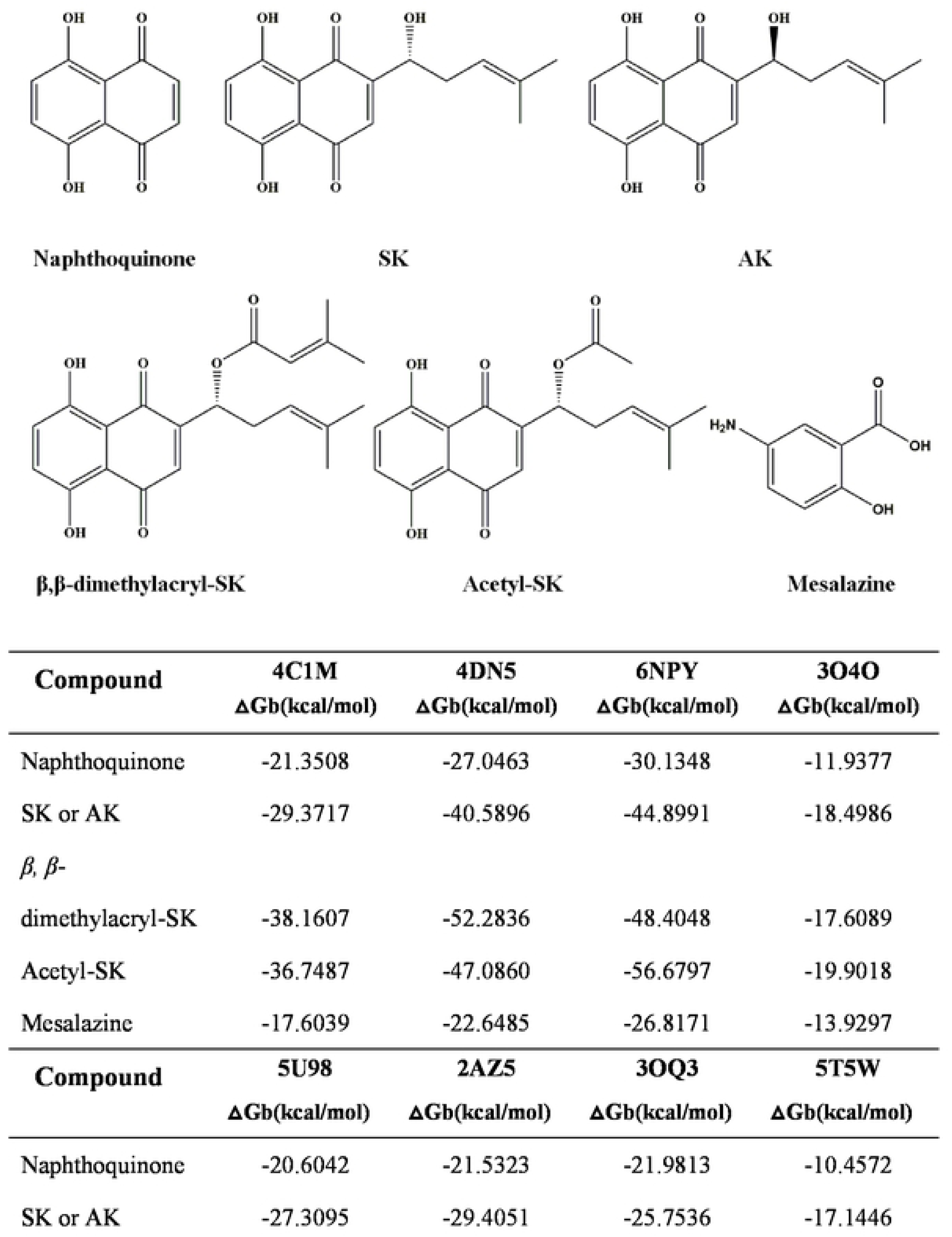
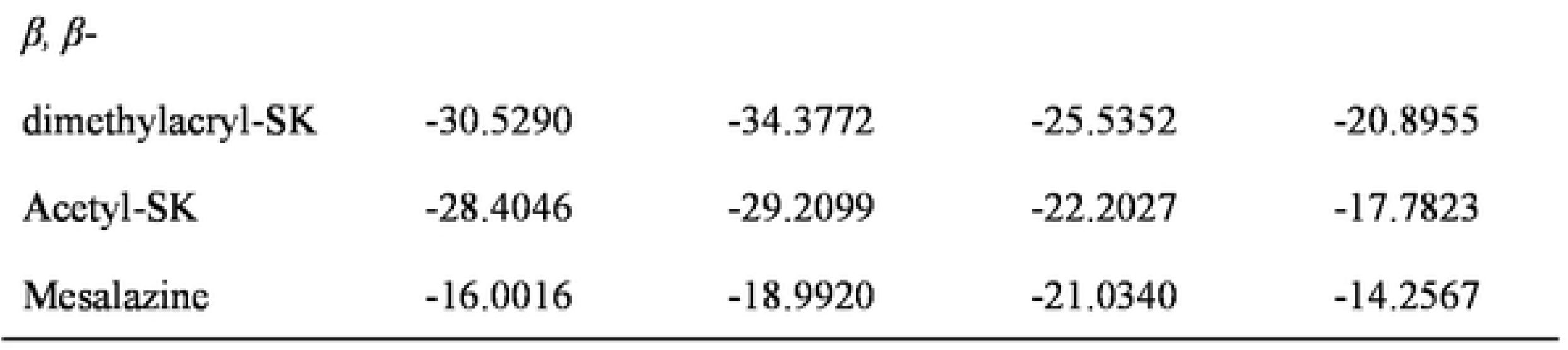
Binding energy of Naphthoquinone, Shikonin or Alkanin, *β, β*-dimethylacrylshikonin, Acetylshikonin and Mesalazine with 4CIM, 4DN5, 6NPY, 3040, 5U98, 2AZ5, 30Q3 and 5T5W.

| Compound | 4C1M<br>$\Delta G_b(\text{kcal/mol})$ | 4DN5<br>$\Delta G_b(\text{kcal/mol})$ | 6NPY<br>$\Delta G_b(\text{kcal/mol})$ | 3O4O<br>$\Delta G_b(\text{kcal/mol})$ |
| --- | --- | --- | --- | --- |
| Naphthoquinone | -21.3508 | -27.0463 | -30.1348 | -11.9377 |
| SK or AK | -29.3717 | -40.5896 | -44.8991 | -18.4986 |
| $\beta$, β-<br>dimethylacryl-SK | -38.1607 | -52.2836 | -48.4048 | -17.6089 |
| Acetyl-SK | -36.7487 | -47.0860 | -56.6797 | -19.9018 |
| Mesalazine | -17.6039 | -22.6485 | -26.8171 | -13.9297 |
| Compound | 5U98<br>$\Delta G_b(\text{kcal/mol})$ | 2AZ5<br>$\Delta G_b(\text{kcal/mol})$ | 3OQ3<br>$\Delta G_b(\text{kcal/mol})$ | 5T5W<br>$\Delta G_b(\text{kcal/mol})$ |
| Naphthoquinone | -20.6042 | -21.5323 | -21.9813 | -10.4572 |
| SK or AK | -27.3095 | -29.4051 | -25.7536 | -17.1446 |
| $\beta$ , $\beta$ - |  |  |  |  |
| dimethylacryl-SK | -30.5290 | -34.3772 | -25.5352 | -20.8955 |
| Acetyl-SK | -28.4046 | -29.2099 | -22.2027 | -17.7823 |
| Mesalazine | -16.0016 | -18.9920 | -21.0340 | -14.2567 |

**S2 Table.** The standard for the disease activity index (DAI) evaluation.

| Loss of weight (%) | Stool property* | Stool occult blood/gross blood stool | score |
| --- | --- | --- | --- |
| 0 | Normal | Normal | 0 |
| 1-5 |  |  | 1 |
| 6-10 | Loose stool | Slight bleeding | 2 |
| 11-15 |  |  | 3 |
| >15 | Liquid stool | Blood in stool | 4 |

\*: Normal, formed stool; Loose stool, a paste, semi-formed stool that does not adhere to the anus; Liquid stool, a watery stool that can adhere to the anus.

**S3 Table.**
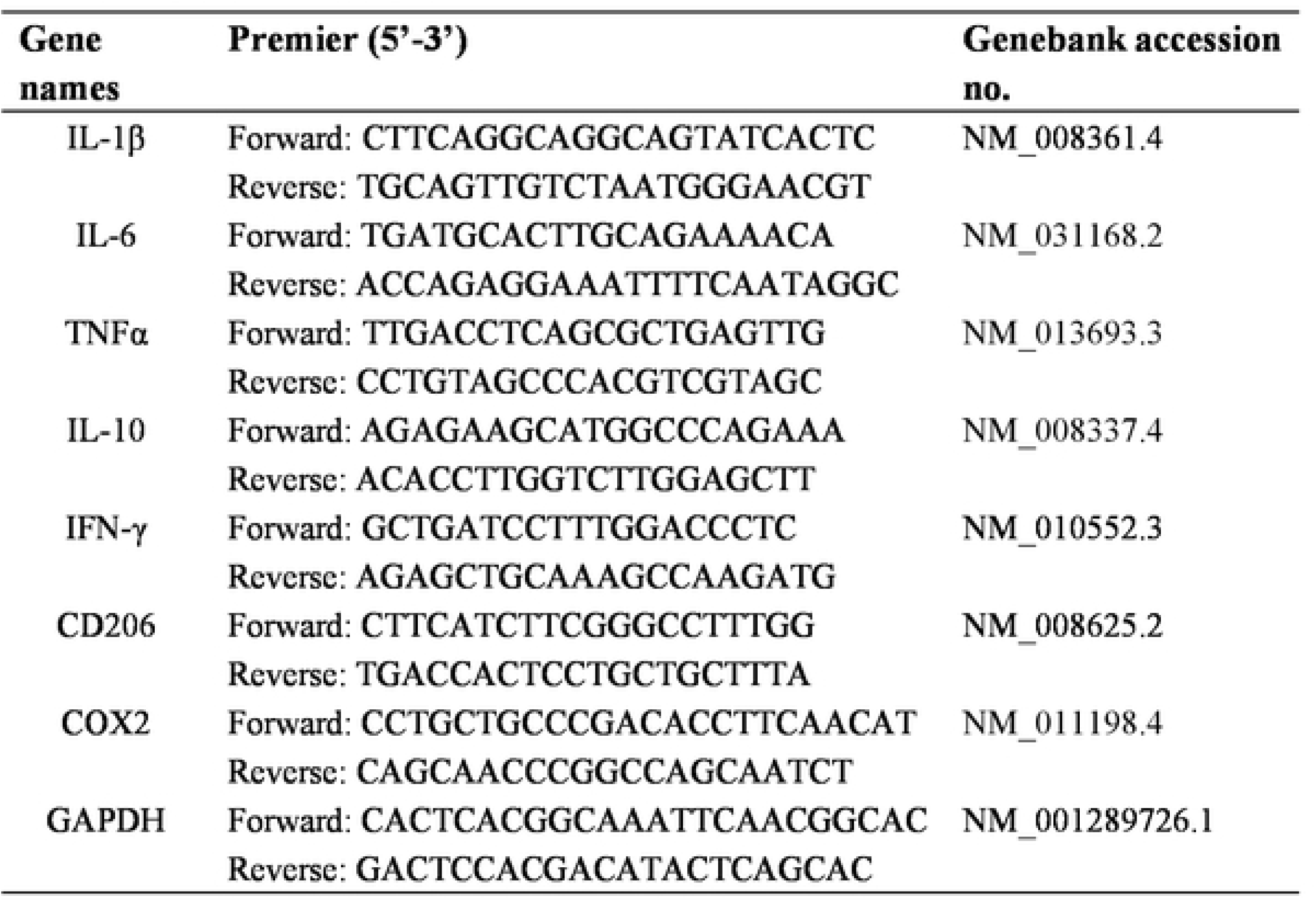
Premier sequences of the genes.

### Representative ^1^HNMR spectra

*5,8-Dihydroxynaphthalenel-,4-dione **(Naphthoquinone,*** PubChem CID: 10141)

Red powder, Mp: 227 °C. ^1^H NMR (600 MHz, CDCl_3_) δ12.41 (s, 2H, -OH), 7.15 (s, 4H, Ar-H). ^13^C NMR (151 MHz, CDC1_3_) δ 172.85 (s, IC, 4C, 5C, 8C), 134.65 (s, 2C, 3C, 6C, 7C), 111.87 (s, 9C, 10C). ESI-TOF, calcd for C_10_H_6_O_4_ ([M +H]^+^), 191.03, found 190.96. Anal. Calcd for C_10_H_6_O_4_: C, 63.16; H, 3.18; O, 33.65; Found: C, 63.08; H, 3.20; O, 33.66.

*(R)-5,8-Dihydroxy-2-(l-hydroxy-4-methylpent-3-en-1-yl)naphthalene-J, 4-dione*

***(Shikonin,*** PubChem CID: 479503)

Red powder, Mp: 147-148 °C. ^1^H NMR (600 MHz, CDC1_3_) δ 12.60 (s, 1H, -OH), 12.49 (s, 1H, -OH), 7.20 (t, J = 7.2 Hz, 2H, Ar-H), 7.18 − 7.17 (m, 1H, -C-CH=C), 5.21 (dd, J = 8.0, 6.9 Hz, 1H, -OH), 4.93 − 4.90 (m, 1H, -C-CH=C), 4.12 (dd, J = 14.3, 7.1 Hz, 1H, -O-CH), 2.67 − 2.62 (m, 1H, -CH_2_), 2.39 − 2.33 (m, 1H, -CH_2_), 1.76 (s, 3H, -CH_3_), 1.66 (s, 3H, -CH_3_). ^13^C NMR (151 MHz, CDCl_3_) δ 180.66 (s, SC), 179.86 (s, SC), 165.48 (s, 4C), 164.87 (s, 1C), 151.44 (s, 7C), 137.48 (s, 6C), 132.41 (s, 14C), 132.30 (s, 2C), 131.87 (s, 3C), 118.44 (s,13C), 112.03 (s, 9C), 11155 (s, 10C), 68.35 (s, l1C), 35.68 (s, 12C), 25.97 (s, 15C), 18.10 (s, 16C). ESI-TOF, calcd for C_16_H_16_O_5_ ([M+H]^+^), 289.10, found 288.78. Anal. Calcd for C_16_H _16_O_5_: C, 66.66; H, 5.59; 0, 27.75; Found: C, 66.65; H, 5.58; 0, 27.72.

*(S)-5,8-Dihydroxy-2-(1-hydroxy-4-methylpent-3-en-l-yl)naphthalene-l,4-dione*

***(Alkanin,*** PubChem CJD: *32465)*

Red powder, Mp: 149 °C. ^1^H NMR (600 MHz, CDCI_3_) δ 12.60 (d, J = 4.8 Hz, 1H, -OH), 12.50 (d, J = 5.0 Hz, 1H, -OH), 7.21 (t, J = 7.2 Hz, 2H, Ar-H), 7.19 − 7.17 (m, 1H, -C-CH=C), 5.21 (ddd, J = 8.1, 2.7, 1.3 Hz, 2H, -OH, -O-CH), 4.92 (dd, J = 7.8, 4.3 Hz, 1H, -C-CH=C), 2.68 − 2.63 (m, 1H, -CH_2_), 2.36 (dd, J = 14.8, 7.6 Hz, 1H, -CH_2_), 1.76 (s, 3H, -CH_3_), 1.66 (s, 3H, -CH_3_). ^13^C NMR (151 MHz, CDC1_3_) δ 180.66 (s, 8C), 179.86 (s, SC), 165.48 (s, 4C), 164.87 (s, 1C), 151.44 (s, 7C), 137.48 (s, 6C), 132.41 (s, 14C), 132.30 (s, 2C), 131.87 (s, 3C), I18.44 (s, 13C), 112.03 (s, 9C), 111.55 (s, 10C), 68.35 (s, 11C), 35.68 (s, 12C), 25.97 (s, 15C), 18.10 (s, 16C). ESI-TOF, calcd for C_1_6Hi60 _5_ ((M+H]+), 289.10, found 288.78. Anal. Calcd for C_1_6H i60 S: C, 66.66; **H,** 5.59; O, 27.75; Found: C, 66.67; H, 5.62; O, 27.71.

*(R)-1-(5,8-Dihydroxy-J,4-dioxo-J,4-dihydronaphthalen-2-y/)-4-methylpent-3-en-1-yl3-methylbut-2-enoate(β, β**-dimethylacrylshikonin,*** PubChem CID: 479499)

Red powder, Mp 103-105 °C. **^1^H** NMR (600 MHz, CDC1_3_) δ 12.60 (s, 1H, -OH), 12.44 (s, 1H, -OH), 7.18 (s, 1H, -C-C H = C), 7.00 − 6.97 (m, 2H, Ar-H), 6.03 (ddd, J = 18.2, 7.3, 4.0 Hz, 1H, -C-CH=C), 5.79 − 5.77 (m, 1H, -O-CH-), 5.15 (t, J = 7.3 Hz, 1H, -C-CH=C), 2.63 (dt, J = 11.7, 6.6 Hz, 1H, -CHr), 2.48 (dt, J = 14.9, 7.3 Hz, 1H, -CH_2_-), 2.15 (d, J = 10.1 Hz, 3 H, -CH_3_), 1.94 (d, J = 0.8 Hz, 3H, -CH_3_), 1.69 (s, 3H, -CH_3_), 1.58 (s, 3H, -C H_3_). ^13^C NMR (151 MHz, CDC1_3_) δ 179.03 (s, 5C), 177.53 (s, 5C), 166.79 (s, 17C), 166.25 (s, 4C), 165.27 (s, IC), 159.01 (s, 19C), 149.05 (s, 7C), 135.86(s, 6C), 132.60 (s, 14C), 132.4S(s,2C), 131.60 (s, 3C), 118.01 (s, 13C), 115.26 (s, 18C), 111.87 (s, 9C), 111.60 (s, 10C), 68.63 (s, 11C), 32.90 (s, 12C), 27.59 (s, 21C), 25.77 (s, 16C), 20.38 (s, 20C), 17.97 (s, 15C). ESI-TOF, calcd for C_21_H_22_O_6_([M+H]^+^), 371.1 4, found 370.10. Anal. Calcd for C_21_H_22_O_6_: C, 68.09; H, 5.99; O, 25.92; Found: C, 68.09; H, 5.98; O, 25.91

*(R)-1-(5,8-Dihydroxy-I.4-dioxo-J,4-dihydronaphthalen-2-yl)-4-methylpent-3-en-l-ylacetate (Acetylshikonin,* PubChem CID: 479501)

Red powder, Mp: 86 °C. ^1^H NMR (600 MHz, CDC1_3_) δ 12.59 (s, 1H, -OH), 12.43 (s, 1H, -OH), 7.19 (s, 2H, Ar-H), 6.99 (s, 1H, -C-CH=C), 6.02 (dd, J = 7.2, 4.6 Hz, 1H, -O-CH-), 5.12 (t, J = 7.3 Hz, 1H, -C-CH=C), 2.65 − 2.60 (m, 2H, -CH_2_-), 2.14 (s, 3H, -CH_3_), l.69 (s, 3H, -CH_3_), 1.58 (s, 3H, -CH_3_). ^13^C NMR(151 MHz, CDC1_3_) δ 178.23 (s,8C), 176.71 (s, 5C), 169.79 (s, 17C), 167.48 (s, 4C), 166.96 (s, 1C), 148.22 (s, 7C), 136.14 (s, 6C), 132.90 (s, 14C), 132.74 (s, 2C), 131.47 (s, 3C), 117.67 (s, 13C), 111.83 (s, 9C), 111.57 (s, 10C), 69.53 (s, 11C), 32.83 (s, 12C), 25.78 (s, 16C), 21.00 (s, 18C), 17.95 (s, 15C). ESI-TOF, calcd for C_18_H_18_O _6_ ([M+H]^+^), 33 1.34, found 331.12. Anal. Calcd for C_18_H_18_O_6_: C, 65.45; H, 5.49; 0, 29.06; Found: C, 65.40; H, 558; 0, 29 03.

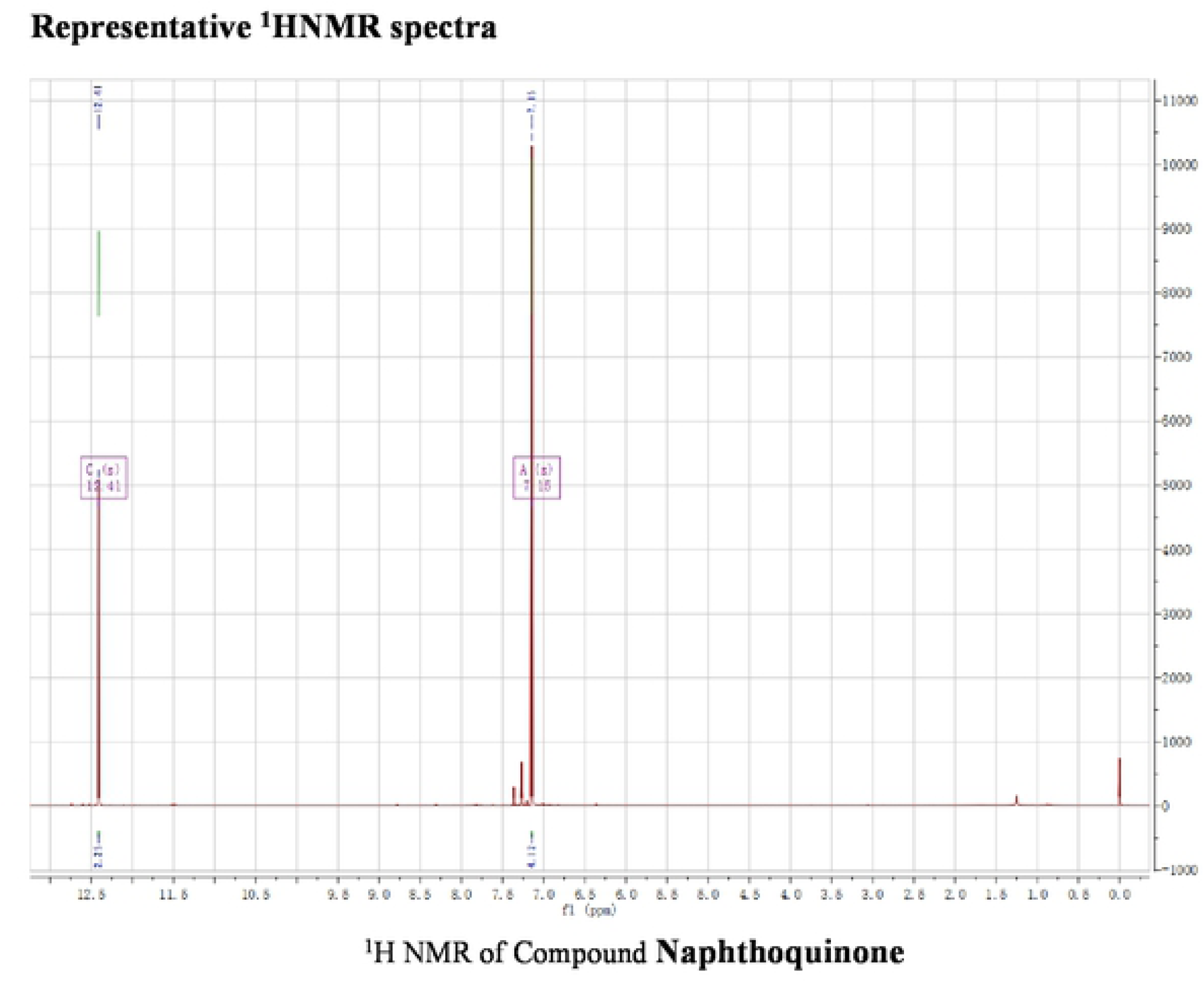
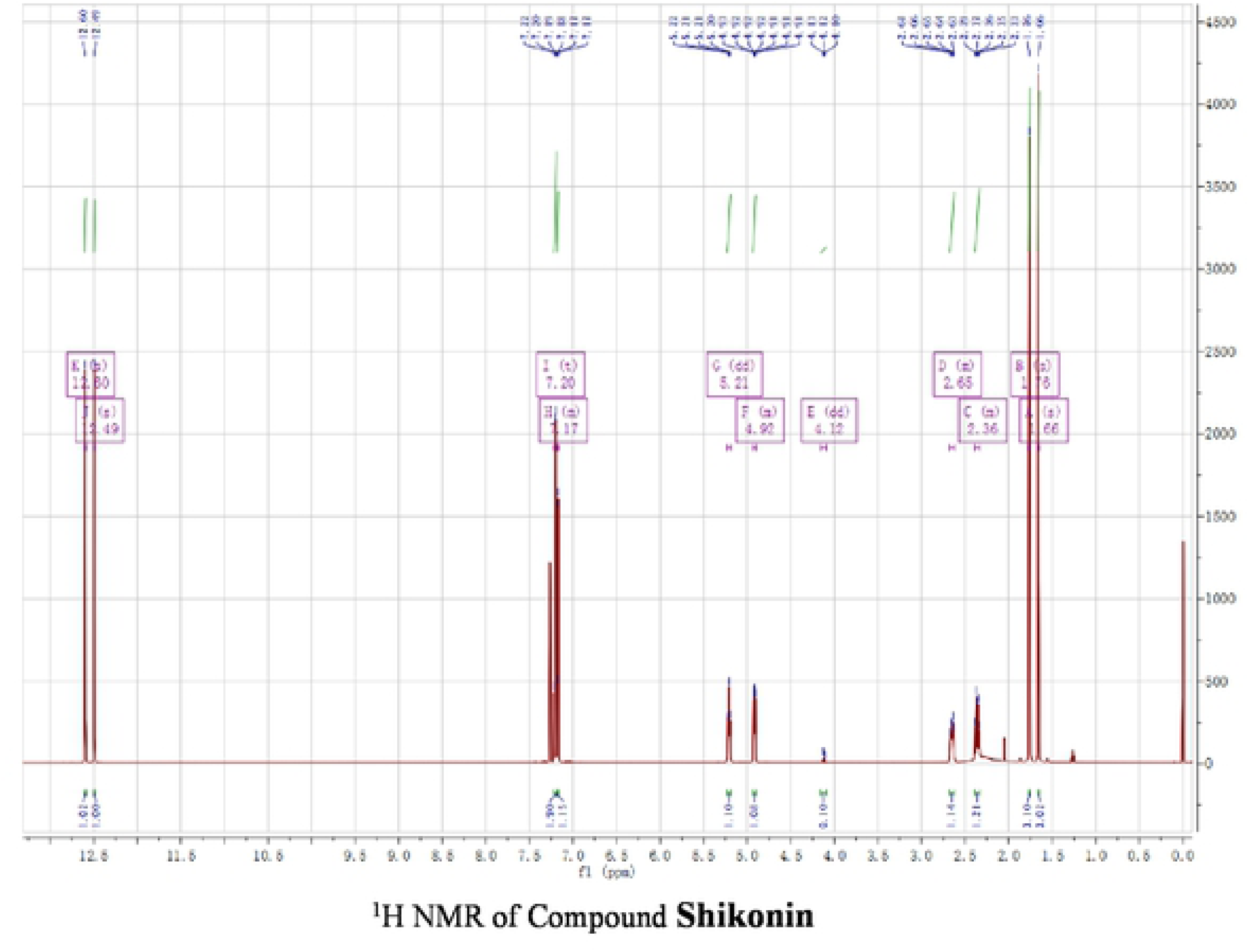
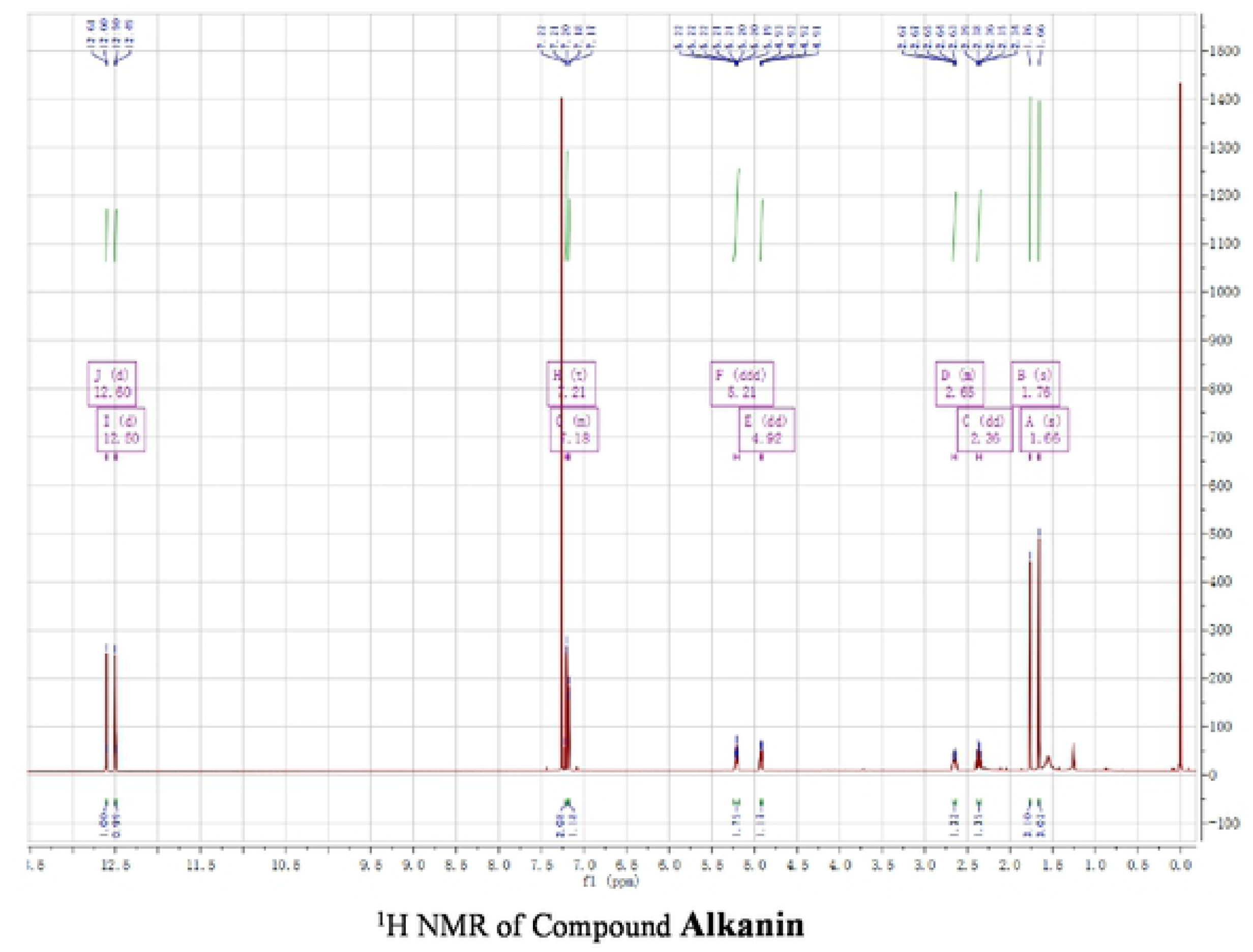
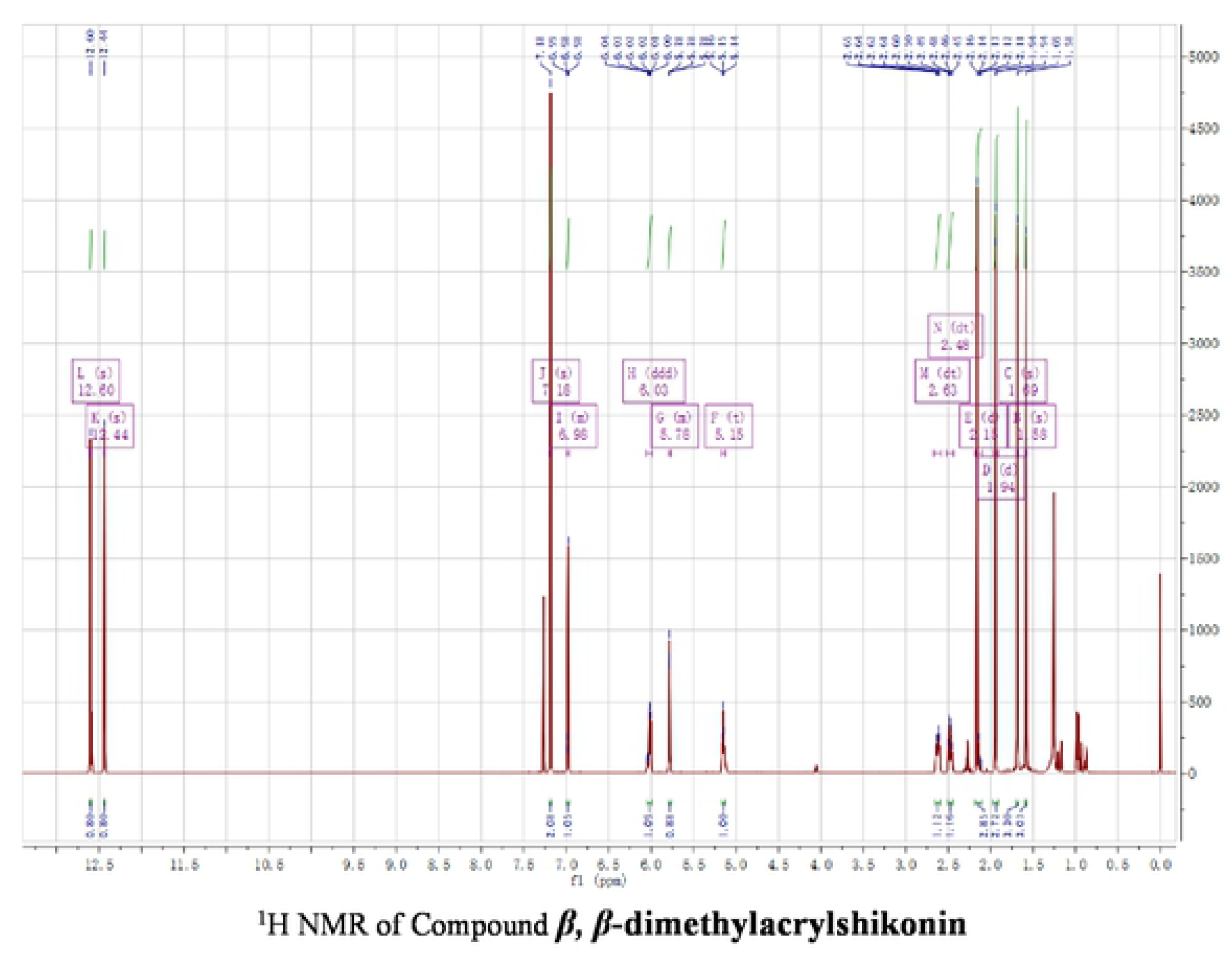
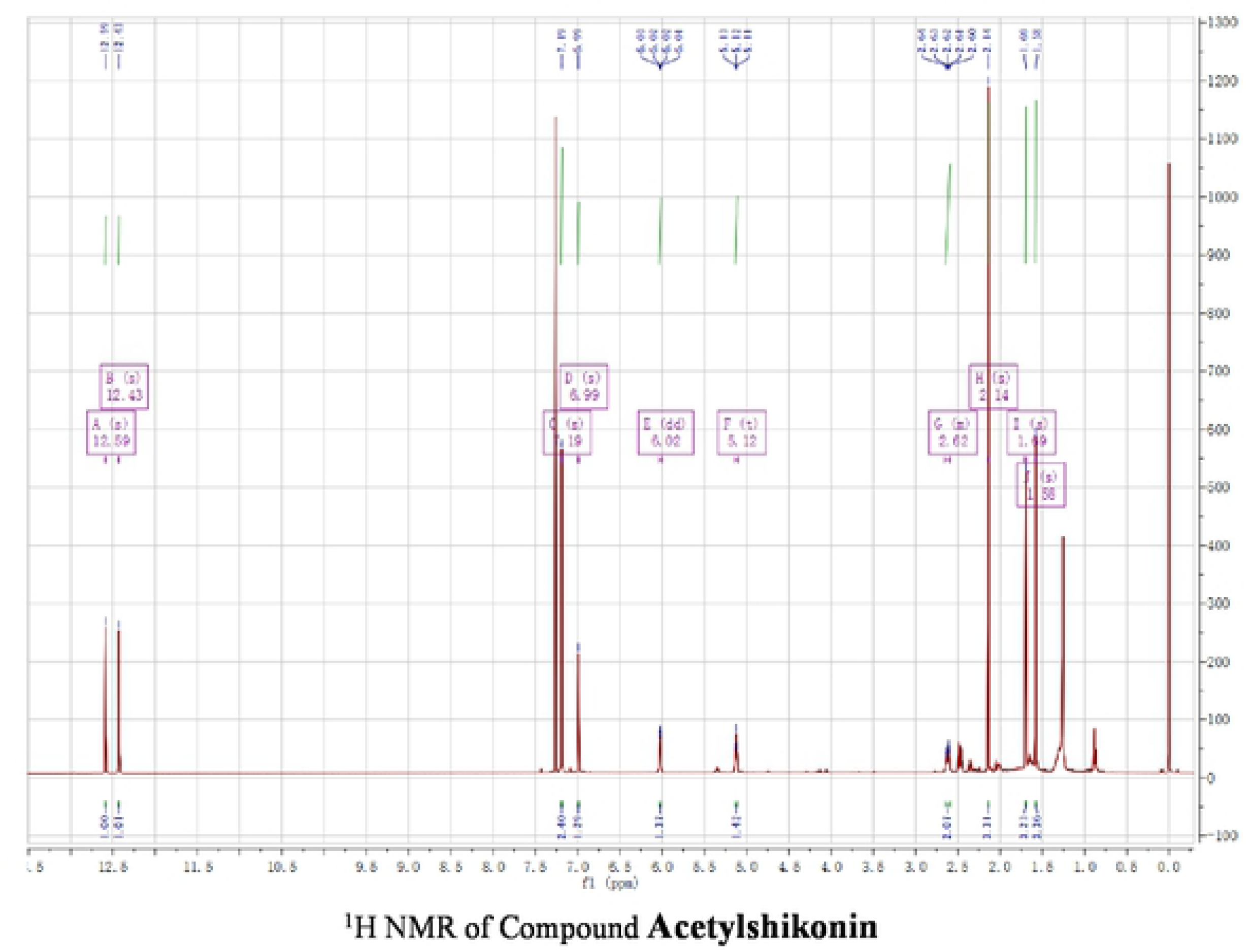
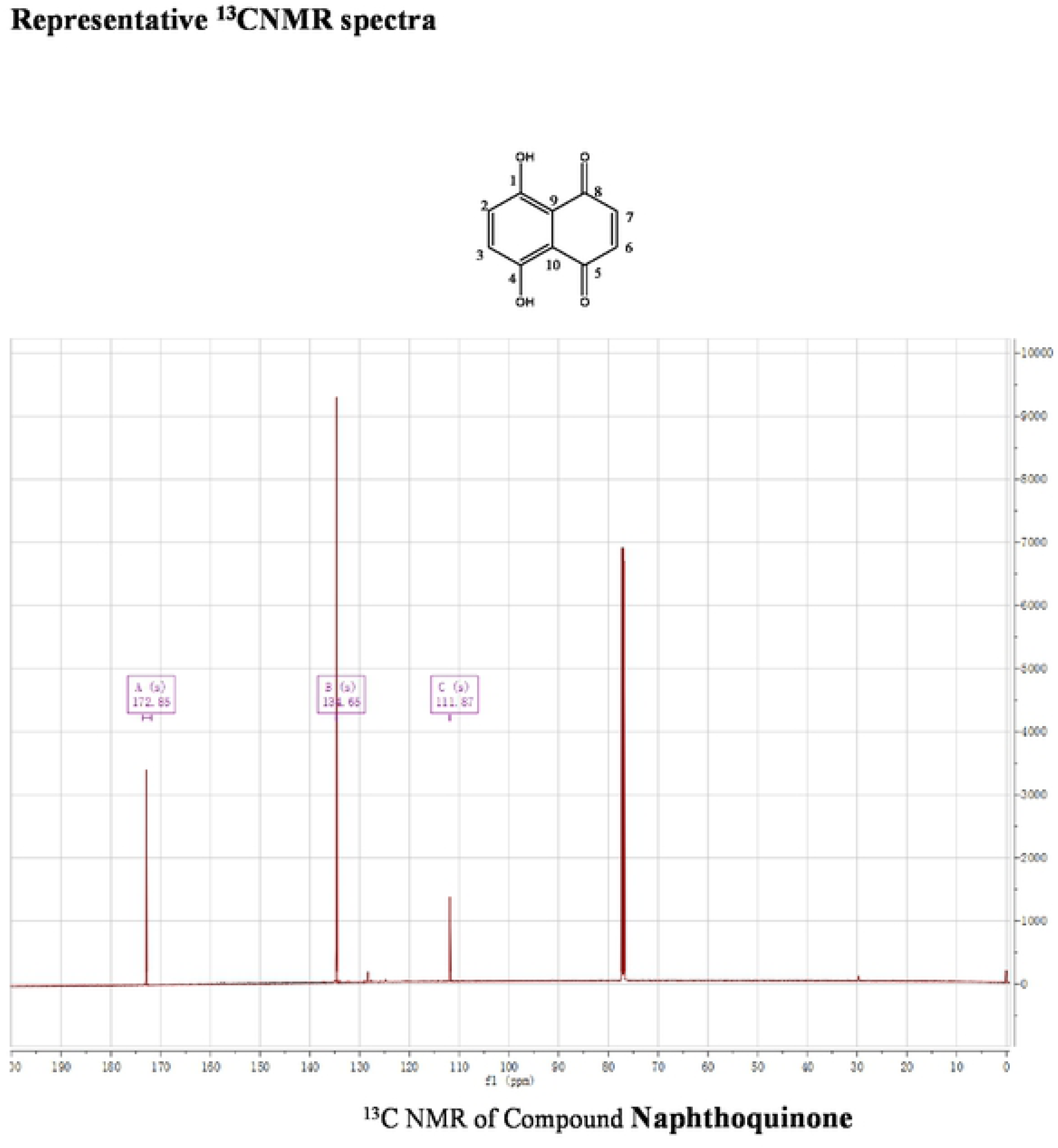
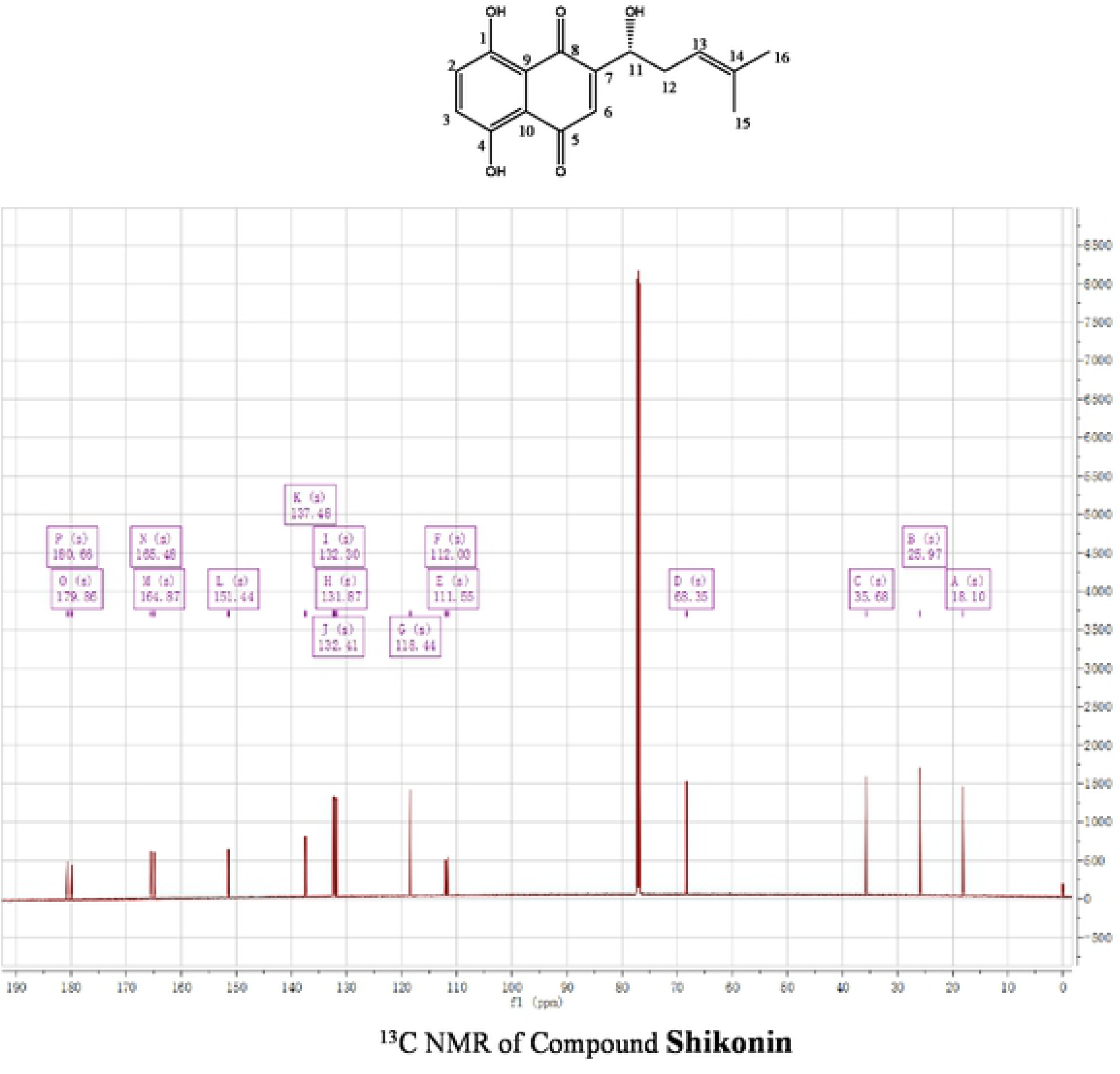
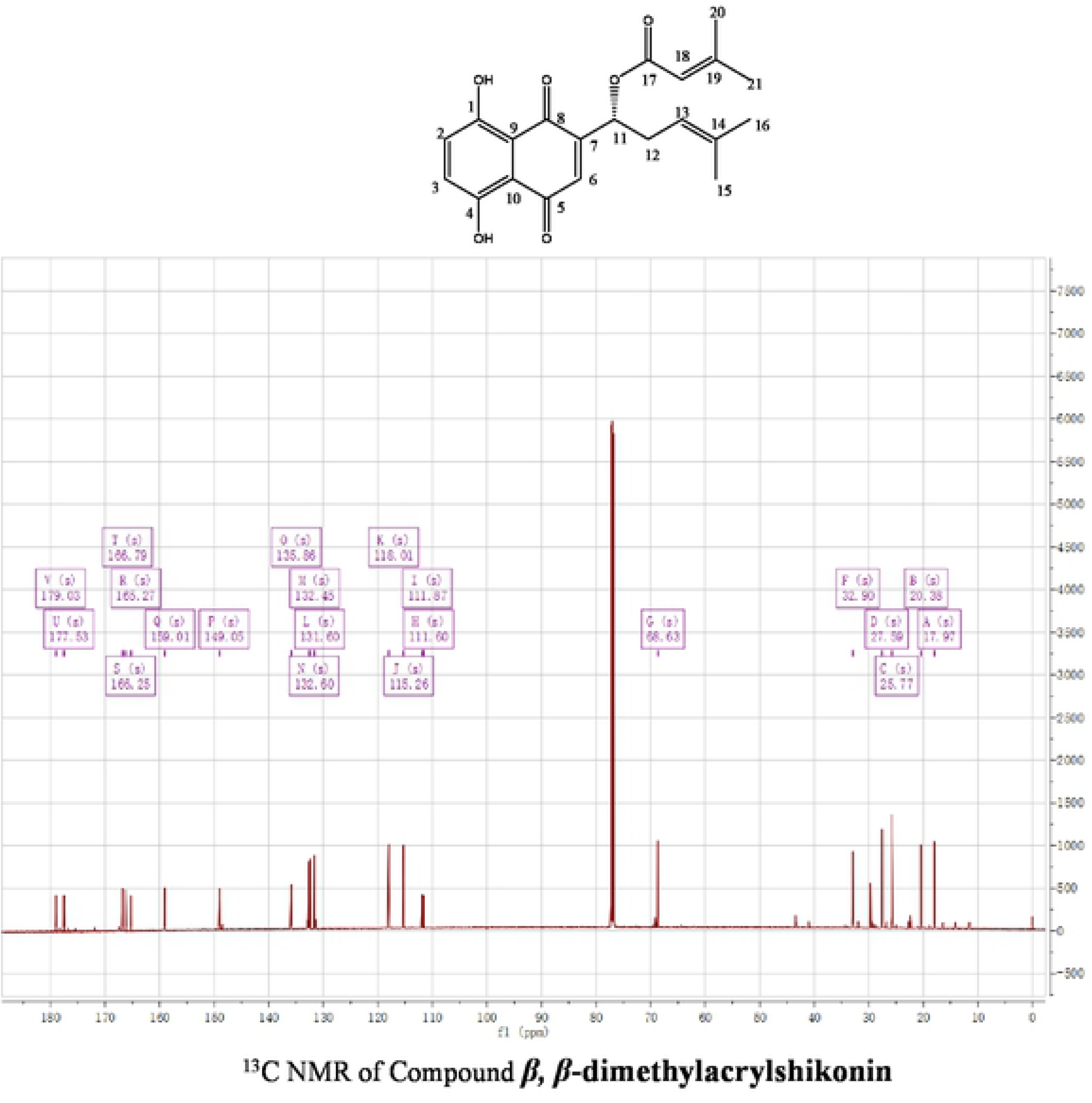
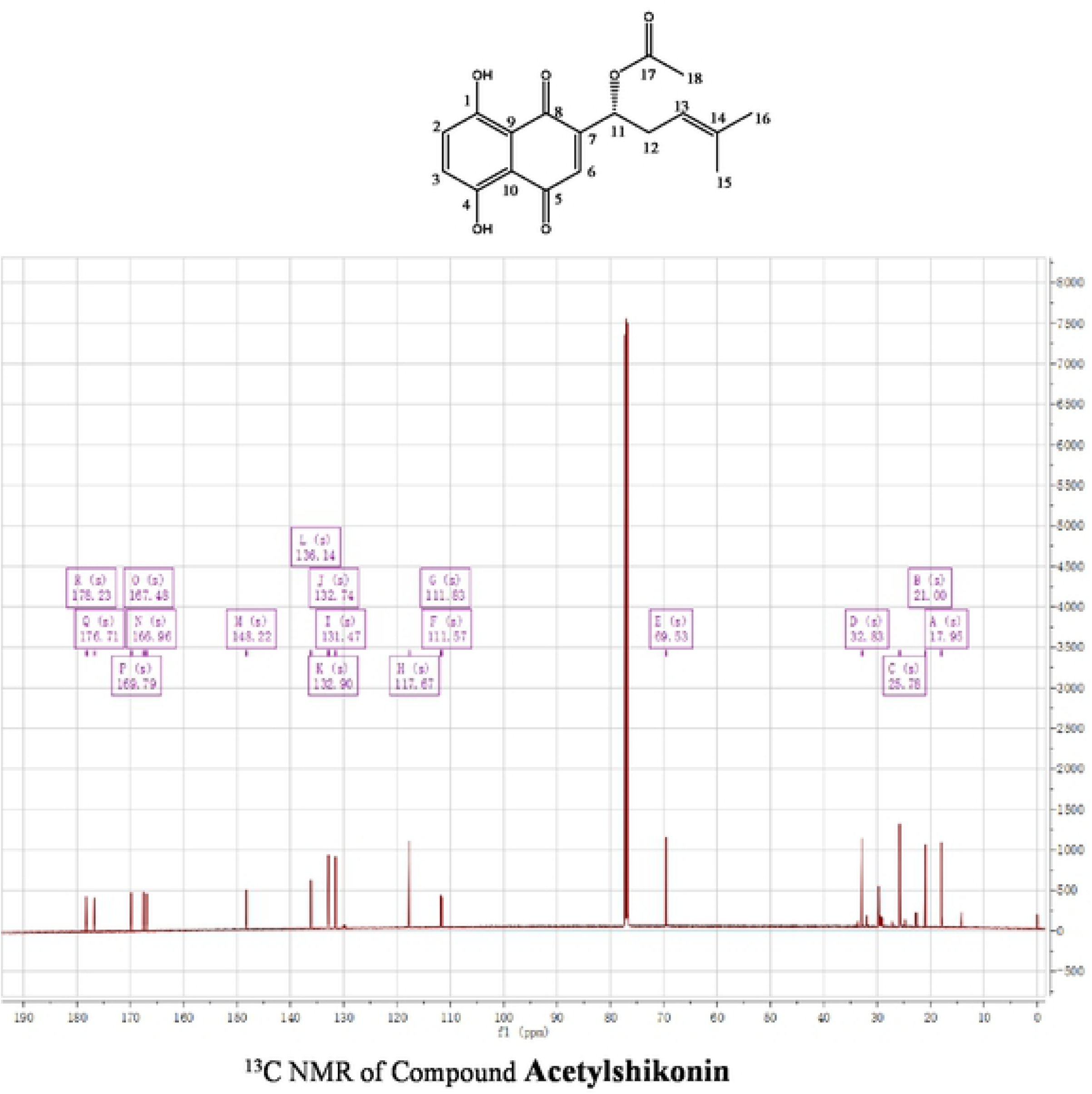

---

**S1 Fig.**
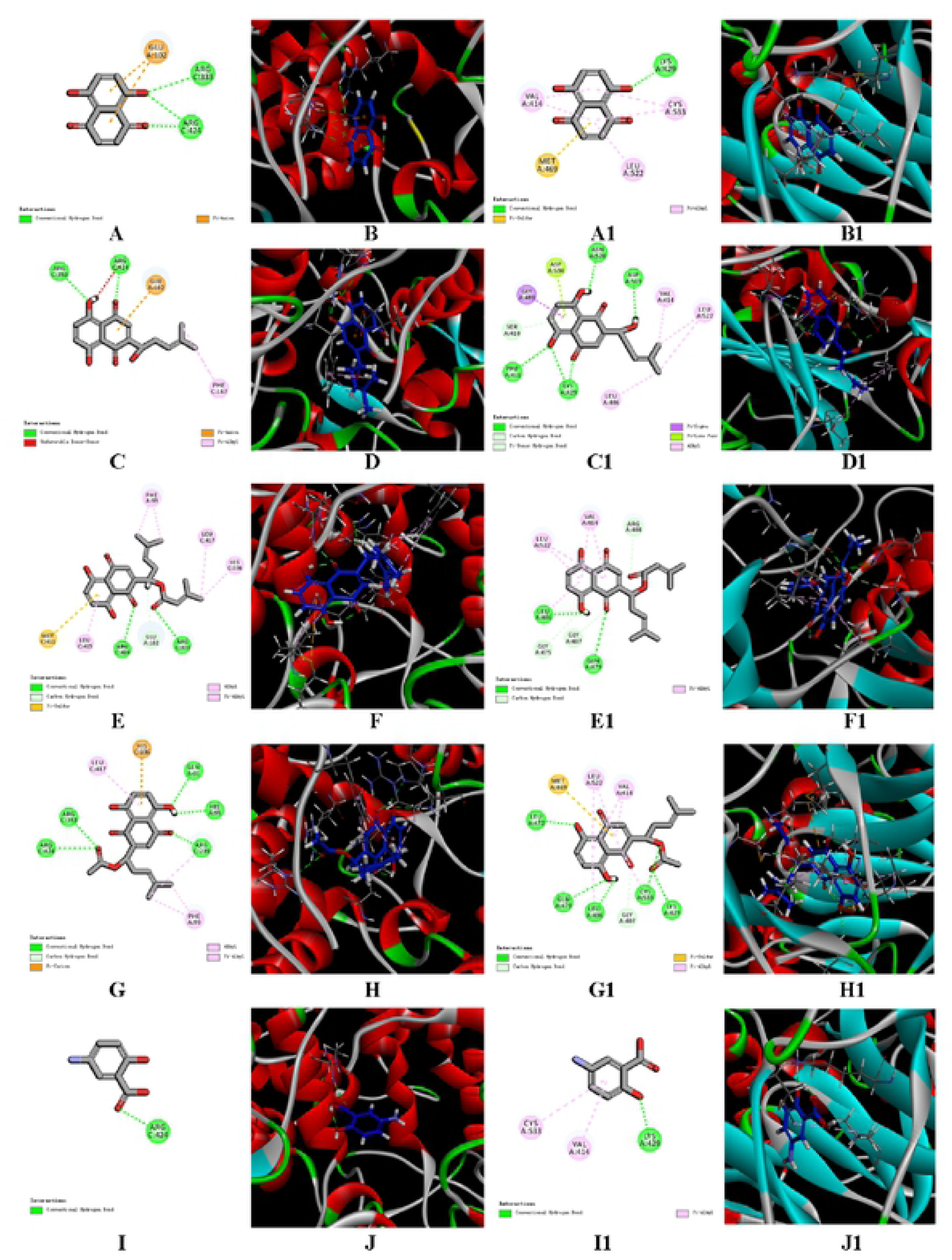
Two and Three Dimensional Molecular Docking Model of Compound Mesalazine, Shikonin and its derivatives with MPO (Entry 4C IM in the Protein Data Bank in the right half part) and with NF-KB (Entry 4DNS in the Protein Data Bank in the left half part).

**S2 Fig.**
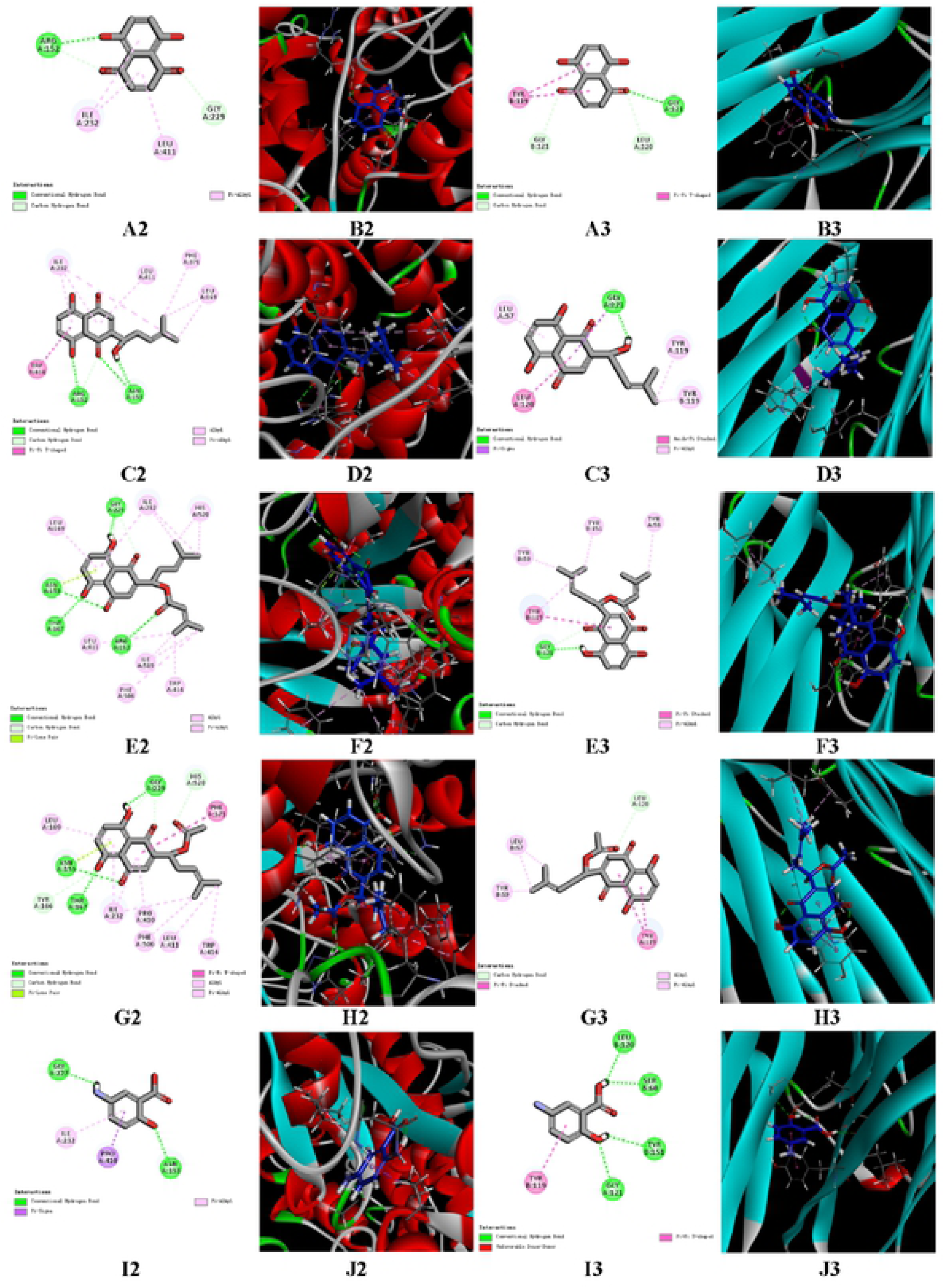
Two and Three Dimensional Molecular Docking Model of Compound Mesalazine, Shikonin and its derivatives with NLRP3 (Entry 6NPY in the Protein Data Bank in the right half part) and with TNF-a (Entry 2AZ5 in the Protein Data Bank in the left half part).

**S3 Fig.**
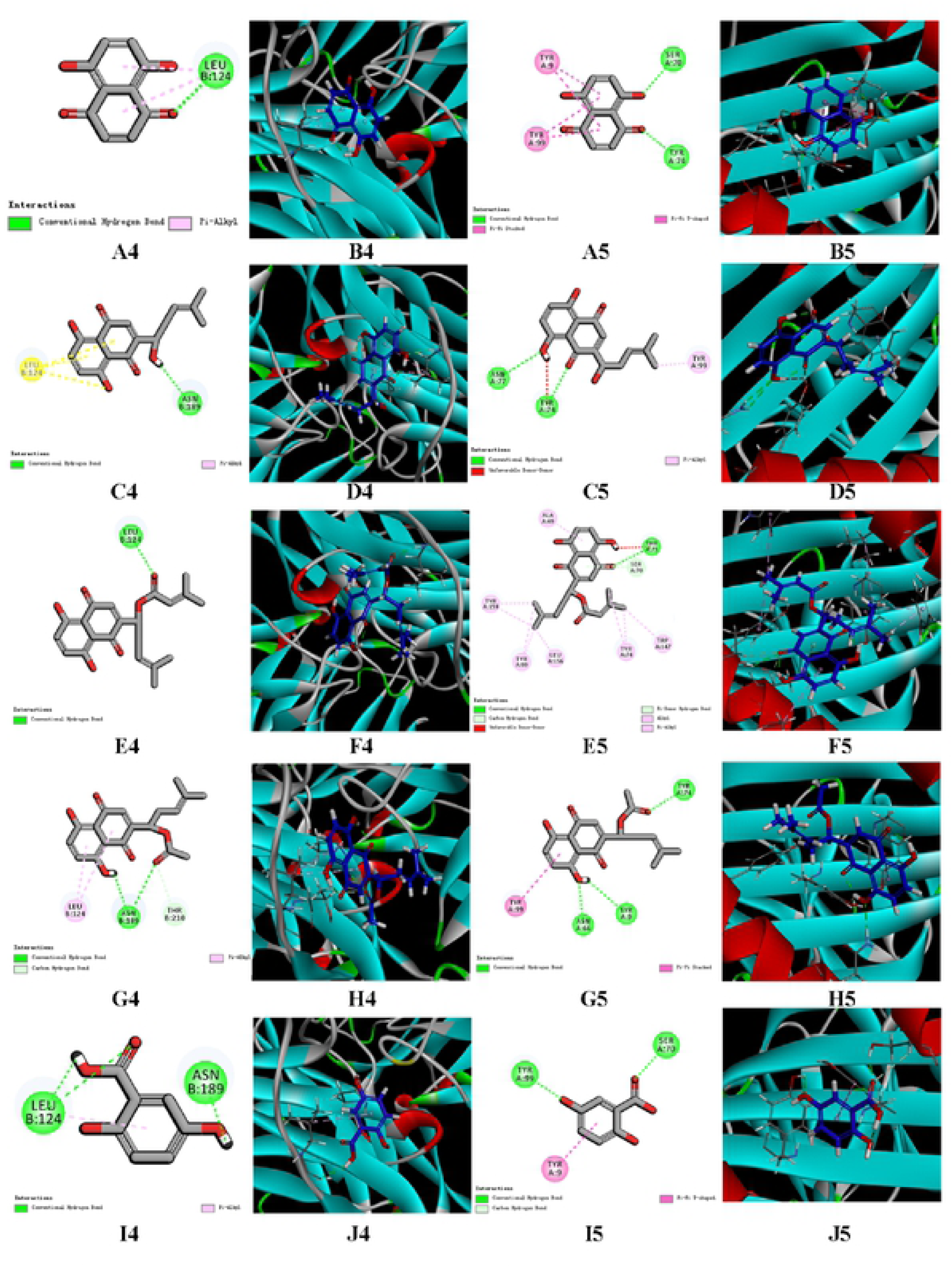
Two and Three Dimensional Molecular Docking Model of Compound Mesalazine, Shikonin and its derivatives with IL-IP (Entry 3040 in the Protein Data Bank in the right half part) and with fL-6 (Entry 5U98 in the Protein Data Bank in the left half part).

**S4 Fig.**
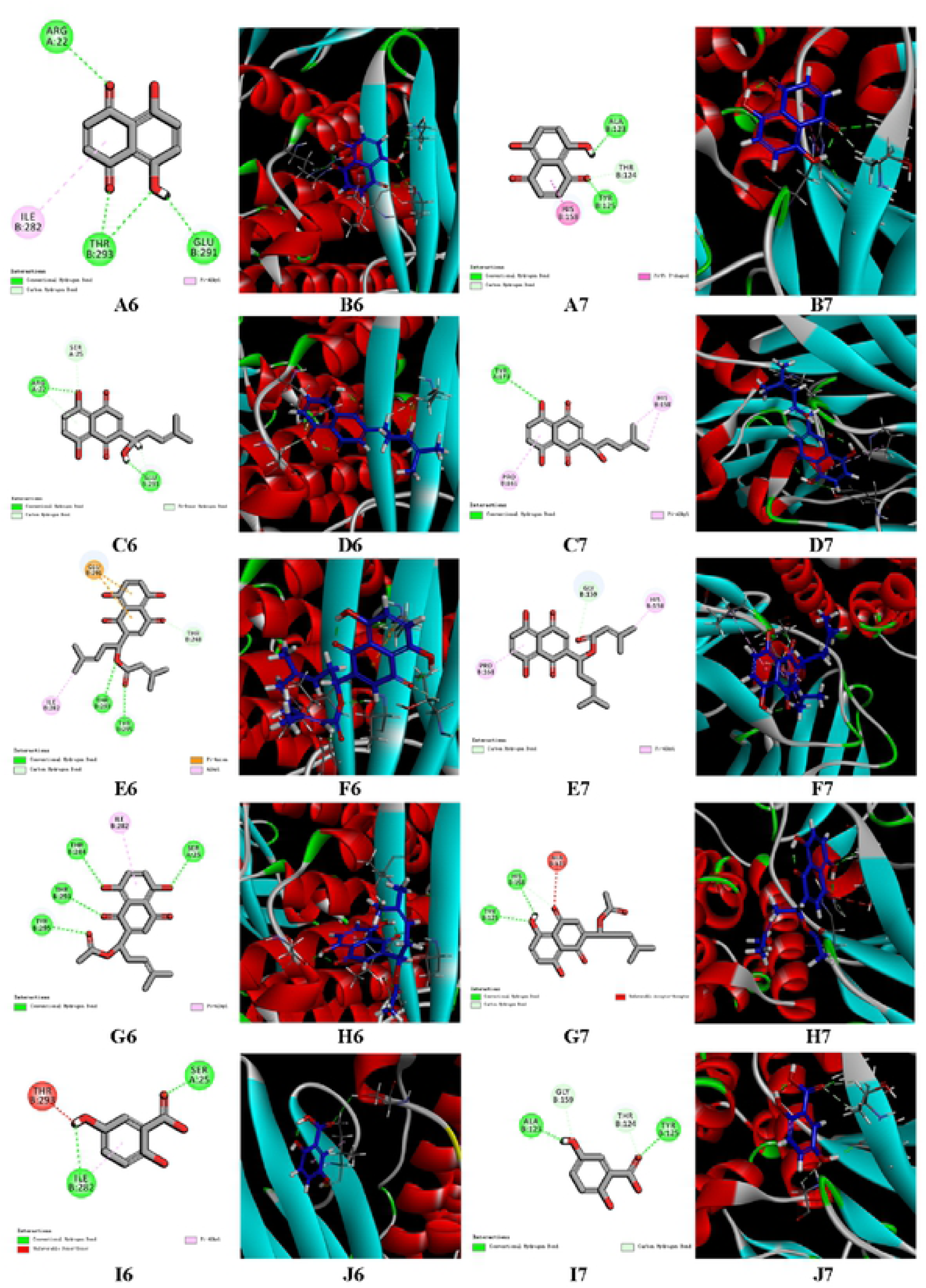
Two and Three Dimensional Molecular Docking Model of Compound Mesalazine, Shikonin and its derivatives with IFNy (Entry 30Q3 in the Protein Data Bank in the right half part) and with IL-IO (Entry STSW in the Protein Data Bank in the left half part).

